# Modulation of β-Catenin is important to promote WNT expression in macrophages and mitigate intestinal injury

**DOI:** 10.1101/2024.09.21.614209

**Authors:** Rishi Man Chugh, Payel Bhanja, Ryan Zitter, Sumedha Gunewardena, Rajeev Badkul, Subhrajit Saha

## Abstract

Macrophages are the major source of WNT ligands. Macrophage-derived WNT is one of the most potent regenerative signals to mitigate intestinal injury. However, regulation of WNT expression in macrophages has not been studied. In the present study, we discovered that activation of canonical β-Catenin suppresses WNT expression in macrophages. Our CHIP-seq and validation study demonstrated the involvement of β-Catenin in the transcriptional regulation of WNT expression. Genetic and pharmacological approaches to de-stabilize/inactivate β-Catenin induce WNT expression in macrophages. Extracellular vesicles (EVs) are a major career of WNT ligands. Transfusion of EVs from pre-conditioned WNT-enriched macrophages demonstrated significant regenerative benefit over native macrophage-derived EVs to mitigate radiation-induced intestinal injury. Transfusion of WNT-enriched EVs also reduces DSS-induced colitis. Our study provides substantial evidence to consider that macrophage-targeted modulation of canonical WNT signaling to induce WNT expression followed by treatment with WNT-enriched EVs can be a lead therapy against intestinal injury..

**SUMMARY:** Activation of β-Catenin suppresses WNT expression in macrophages. Macrophage-targeted pharmacological modulation of canonical WNT signaling followed by adoptive transfer mitigate radiation injury in intestine. EVs from these preconditioned macrophages mitigate chemical or radiation induced intestinal injury.

## INTRODUCTION

Macrophages are one of the major sources of morphogenic WNT ^1^. We have previously reported that macrophages play an important role in intestinal epithelial regeneration ^2^. Our study demonstrated that macrophage derived WNTs may not be crucial for intestinal homeostasis, but critical to promote the repair process against radiation injury. However, the mechanism behind the regulation of WNT expression in macrophages has not been studied.

Radiosensitivity of intestinal epithelium is a major limiting factor for effective radiotherapy to gastrointestinal cancer such as pancreatic cancer or liver cancer. Radiation induced loss of intestinal stem cells (ISC) ^3,4^ impairs epithelial regeneration resulting damage in mucosal barrier. Large dose of radiation may cause irreversible loss of mucosal integrity resulting systemic influx of bacterial pathogens, sepsis, and death ^5,6^. Moreover, victims of radiation accidents or nuclear terrorism are also at high risk for lethal consequences due to radiation induced mucosal damage. So far there are no Food and Drug Administration (FDA) approved agents available to mitigate radiation-induced intestinal injury ^7^. Sepsis-like inflammatory states can also occur in acute colitis with significant loss of ISCs. Repair and regeneration of mucosal epithelium is important to mitigate colitis ^8^. Intestinal epithelial homoeostasis depends on the signaling crosstalk between the ISC and the surrounding niche, including the intestinal subepithelial myofibroblasts, endothelial cells and macrophages. The cells in the ISC niche provide critical growth factor/signals for ISC regeneration and intestinal homoeostasis ^9^.

Considering the importance of WNT signaling in ISC homeostasis and repair several studies have been performed about source, function, and characterization of physiological WNT. There are 19 different isoforms of WNT ligands ^10^. Both epithelial and stromal cells express and release WNT ligands ^10^. Different cell types of intestinal stroma including endothelial cells, macrophages, neurons, fibroblasts and myofibroblasts could produce a cocktail of redundant WNT ligands maintaining intestinal homoeostasis *in vivo*. However, inhibition of WNT release from intestinal myofibroblast ^11^ or from intestinal epithelium ^12^ does not affect intestinal homoeostasis. Moreover, it was noted that global pharmacologic but not epithelial-specific inhibition of *Porcn* significantly increased the radio-sensitivity of intestine ^12^. These observations clearly suggest the importance of stromal cell derived WNTs in intestinal homoeostasis and regeneration. However, regulation of Wnt expression remain poorly understood. Our previous report demonstrated that macrophage is an important source of WNT (2). One recent study demonstrated that lung macrophages promote epithelial proliferation and repair post-injury through a Trefoil factor 2-dependent mechanism that leads to the upregulation of Wnts 4 and 16 ^13^. Elucidation of macrophage derived WNT expression is important to develop macrophage derived WNT as regenerative therapy.

The present manuscript examined the transcriptional regulation of macrophage derived WNT expression. Using both genetically modified mice model and adopting pharmacological approach we discovered that canonical WNT signaling has a negative feedback regulatory loop on WNT expression in macrophages. Macrophage specific deletion of β-Catenin improves the WNT expression several folds compared to control and minimize the intestinal epithelial radiosensitivity. Chip-seq analysis suggested physical association of β-Catenin with WNT promoters to suppress WNT expression. Finally, we have shown that macrophages with perturbed WNT signaling can be a better candidate for adoptive cell therapy to mitigate radiation induced intestinal injury by delivering optimum level of morphogenic WNT to injured epithelium with activation of canonical WNT signaling. Moreover, EVs derived from this WNT enriched macrophages can reduce radiation induced intestinal injury or DSS induced acute colitis in mice.

## RESULTS

### Pharmacological inhibition of canonical WNT signaling in macrophages induces the macrophage derived WNT expression

Macrophages are one of the major sources of WNT ligands ^3,14^. Canonical WNT ligands primarily activate WNT/β-Catenin signaling and stabilize and translocate cytosolic β-Catenin to the nucleus which eventually modulates multiple gene programming related to growth factors, cell differentiation factors etc. ^15–17^. We first examined whether activation of canonical WNT signaling further promotes WNT expression. LiCl an agonist of canonical Wnt signaling ^18^ was used to trigger β-Catenin activation in ex vivo cultured bone marrow macrophages (BMMɸ) from C57BL/6 mice. Conditioned medium (CM) from BMMɸ treated with/without LiCl (10mM, for 4 hours at 37^0^C) was then tested for the presence of functional WNT using TCF/LEF luciferase (TOPflash) assay ^19^. Surprisingly, the CM from LiCl treated BMMɸ demonstrated significant inhibition of luciferase activity compared to untreated BMMɸ (Figure 1A, Figure S1A). To confirm that inhibition in functional WNT is the result of suppression of genes expressing WNT ligands we performed qPCR using RNA purified from BMMɸ treated with/without LiCl treatment. Significant downregulation of Wnt gene expression was observed with LiCl treatment suggesting a possible inhibitory role of canonical WNT/β-Catenin signaling in macrophage derived WNT expression (Figure 1B).

**Figure 1:**
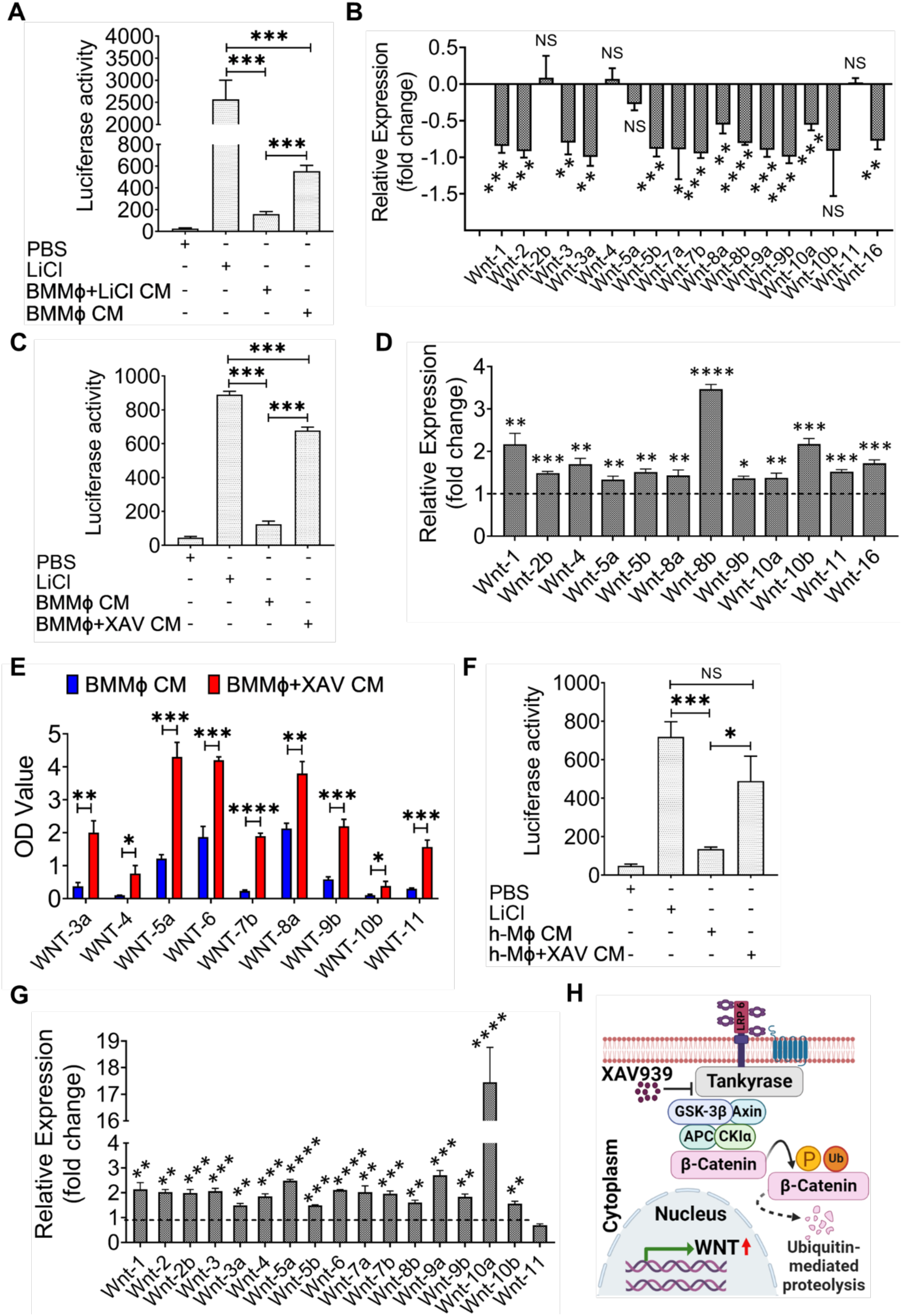
Activation of canonical WNT signaling in BMMɸ inhibits WNT expression in Mɸ: (A) TOPflash assay to determine WNT activity in BMMɸ CM. CM from LiCl-treated BMMɸ showed lower WNT activity compared to untreated BMMɸ CM to. (B) qPCR analysis of RNA from BMMΦ treated with LiCl demonstrated downregulation of Wnt mRNA expression in comparison to the untreated BMMɸ. (C) TOPflash assay using CM from BMMɸ treated with Tankyrase inhibitor XAV939 demonstrated higher WNT activity compared to untreated BMMɸ CM. PBS and LiCl were used as negative and positive control, respectively. (D) qPCR analysis of RNA from XAV939 treated BMMΦ showed significantly higher Wnt expression compared to untreated BMMɸ. (E) ELISA to detect secretory WNTs in BMMɸ conditioned media. (F) TOPflash assay demonstrated higher WNT activity in CM from h-Mɸ treated with XAV939 compared to untreated h-Mɸ CM. (G) qPCR analysis of RNA from XAV939 treated h-Mɸ shows the significantly higher WNT expression compared to untreated h-Mɸ. (H) Schematic representation of canonical WNT signaling with/without pharmacological inhibition of β-Catenin. Data presented as the mean ± SD. (Significant level, *: p<0.05, **: p<0.005, ***: p<0.0005, ****: p<0.00005, NS: Not significant).

To validate this observation, we examined the WNT expression in macrophages treated with Tankyrase inhibitor XAV-939, an inhibitor of canonical WNT signaling pathway ^20,21^. Inhibition of Tankyrase stabilizes AXIN, a key component of the β-Catenin degradation complex, and thereby promotes degradation of β-Catenin and inhibition of canonical WNT signaling ^22^. Ex-vivo culture of mice BMMɸ and human Peripheral Blood Mononuclear Cells (PBMC) derived macrophages was incubated with XAV-939 (10-20μM) for 2 hours.

Degradation of β-Catenin in macrophages in response to XAV-939 was confirmed by immunoblot analysis demonstrating a decrease in the expression of β-Catenin compared to untreated mouse BMMɸ or human macrophages (h-Mɸ) (Figure S2A-C).

To determine the timepoint to detect the optimum level of physiological WNT in BMMɸ CM, BMMɸ was incubated with XAV-939 at 37^0^C for 2 to 4 hours followed by withdrawal of treatment by replacing the media with cell culture media. 24- and 48-hours post-treatment CM from both treated and untreated BMMɸ was then tested for WNT activity using the TOPflash assay. CM collected at 24 hours post treatment demonstrated significantly higher level of WNT activity (Figure 1C, Figure S1B and S3A). qPCR analysis of these BMMɸ derived RNA demonstrated that inhibition of β-Catenin in BMMɸ upregulates the WNT expression compared to untreated cells (Figure 1D). ELISA using these BMMɸ CM demonstrated significantly higher expression of WNT-3a, WNT-4, WNT-5a, WNT-6, WNT-7b, WNT-8a, WNT-9b, WNT-10b and WNT-11 compared to wildtype BMMɸ CM (Figure 1E).

Next, we treated PBMC derived h-Mɸ with XAV-939 following the treatment condition used in mice and tested for WNT activity using the TOPflash assay. CM from XAV-939 treated h-Mɸ shows a significant increase in WNT activity compared to untreated samples (Figure 1F, Figure S1C). qPCR analysis of h-Mɸ derived RNA demonstrated that inhibition of β-Catenin in h-Mɸ upregulates the WNT expression compared to untreated h-Mɸ (Figure 1G).

These data suggest that the pharmacological inhibition of β-Catenin in either mouse BMMɸ or h-Mɸ using a small molecule XAV-939 upregulates the WNT expression (Figure 1 H).

### β-Catenin interacts the WNT Promoter to regulate the transcription of WNT genes

The inhibition of β-Catenin destruction leads to increased levels of β-Catenin, which accumulates in the cytoplasm and then translocate into the nucleus to get involved in vast majority of Wnt/β-Catenin signaling outputs. Since β-Catenin does not possess a DNA binding domain it needs DNA binding partners to bring it to the promoters of its target genes ^23,24^ and β-Catenin acts as the central transcriptional activator. TCF/LEF (hereafter referred to as TCF) transcription factors serve as the main nuclear partners of β-Catenin guiding it to specific DNA loci ^25^. To examine the involvement of β-Catenin in association with TCF in transcriptional regulation of WNT genes a Chip assay was performed with anti-β-Catenin antibody in BMMɸ treated with LiCl. Samples obtained at the end of the Chip were subjected to western blot to determine the presence of β-Catenin and TCF transcription factor (Figure S3B).

Chromatin Immunoprecipitation (Chip) samples were then subjected to DNA sequencing (Figure 2A). Analysis of sequencing data using reference genome by SOAPaligner/SOAP2 (Version: 2.21t) identified the presence of multiple Wnt promoter sequences. The alignment of these Chip DNA samples after sequencing demonstrated Wnt 5a and Wnt 9b promoters have the highest level of confidence scores (Figure 2B). The presence of WNT promoters was further confirmed by qPCR using WNT promoter specific primers, which demonstrated enrichment of multiple WNT promoters in the Chip samples including Wnt3, Wnt4, Wnt5a, Wnt7b, Wnt8a, Wnt9a, Wnt9b, Wnt10b and Wnt11 (Figure S3C). These results further suggested that β-Catenin participates in transcriptional complex consisting of Wnt promoter. To validate the inhibitory role of β-Catenin on WNT promoter activity we developed a recombinant DNA construct expressing luciferase reporter under Wnt 5a and Wnt 9b promoter (Figure 2C). HEK293T cells were then transfected with a respective reporter construct pTA-WNT5a or pTA-WNT9b consisting of Wnt 5a and Wnt 9b promoter or scramble pTA-Luc followed by XAV939 treatment to inhibit β-Catenin activation. Luciferase activity assay demonstrated a significant increase in WNT5a (p<0.0005) or WNT9b (p<0.005) promoter activity in XAV939 treated cells compared to untreated control (Figure 2D, Figure S1D).

**Figure 2:**
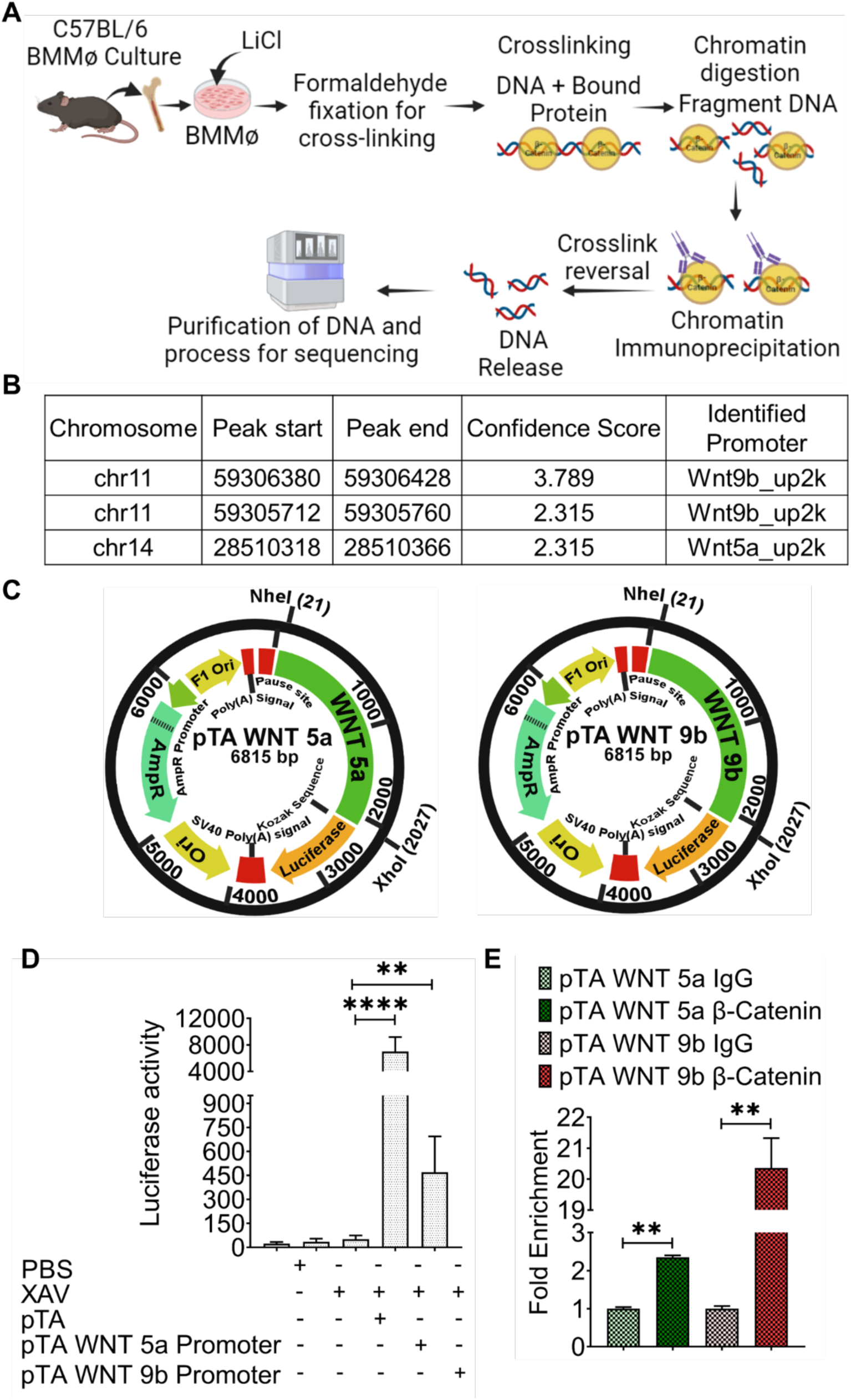
β-Catenin interacts the WNT Promoter to regulate the transcription of WNT genes: (A) Schematic representation of β-Catenin ChIP followed by DNA sequencing in C57BL/6 mouse BMMɸ. (B) ChIP sequence data showed the presence of multiple WNT promoters. WNT9b and WNT5a promoters are top in the list based on confidence score. (C) Construct map of a luciferase reporter system under WNT5a or WNT9b promoter. (D) Luciferase reporter assay to determine WNT5a or WNT9b promoter activity. HEK293T cells transfected with scramble or pTA-WNT 5a or pTA-WNT 9b promoter plasmid followed by detection of luciferase activity. Transfected HEK293T cells treated with XAV939 demonstrated significant increase in luciferase activity (pTA-WNT 5a promoter p<0.00005) and (pTA-WNT 9b promoter p<0.005) compared to scramble plasmid transfected H293T cells. (E) qPCR analysis of β-Catenin CHIP sample from transfected HEK293T cells showed increase in presence of WNT5a promoter 2.35-fold (p<0.005) and WNT9b promoter 20.36-fold (p<0.005) in comparison with their respective IgG control. Data presented as the mean ± SD. (Significant level, **: p<0.005, ****: p<0.00005).

To validate whether β-Catenin physically associated with transcription complex consisting of Wnt 5a and Wnt 9b promoter region, we performed a ChIP assay in HEK293T cells having pTA-WNT5a or pTA-WNT9b construct pre-treated with LiCl to trigger β-Catenin activity. ChIP assay was performed using anti-β-Catenin or rabbit IgG control antibody.

Samples obtained at the end of the Chip assay were analyzed using quantitative PCR with a specific pair of primers for the WNT5a and WNT9b promoters. As demonstrated in Figure 2E, both Wnt5a and Wnt9b promoters were significantly enriched (Wnt5a: 2.35-fold, p<0.005 and Wnt9b: 20.36-fold, p<0.005) in the CHIP sample obtained using immunoprecipitation by the anti-β-Catenin antibody, compared with the normal rabbit IgG control. As expected, the WNTs promoter was not enriched in any one of the negative controls (pTA-WNT5a or pTA-WNT9b). The cells treated with scramble (pTA-Luc) construct followed by immunoprecipitation with the anti-β-Catenin antibody did not show any enrichment of WNT5a and WNT9b promoter through qPCR compared with the normal rabbit IgG control, which suggests that the cells themselves do not have any endogenous WNT promoters. Taken together, these data suggest that β-Catenin physically interacts with WNT5a and WNT9b promoters.

### Deletion of gene expressing β-Catenin (*Ctnnb1)* upregulates macrophage-derived WNT release without any adverse effect on mice health

Transient Pharmacological inhibition of β-Catenin triggered the WNT expression in both mice and human macrophages without altering macrophage survival, characteristics, or function. We next examined whether deletion of gene expressing β-Catenin in macrophages demonstrates similar result. Mice having deletion of β-Catenin restricted to macrophages was achieved by crossing female mice carrying a floxed allele of β-Catenin (*Ctnnb1*) to *Csf1r.iCre* male mice to generate *Csf1r.iCre; Ctnnb1^fl/fl^*mice ^26^. *Csf1r.iCre; Ctnnb1^fl/fl^*mice appeared phenotypically normal, having similar lifespans (followed up to 18 months of age) and body weight (Figure S4A-F) compared to wild type (WT) littermate suggesting macrophage specific deletion of *Ctnnb1* does not influence the overall physiology of the mice.

TOPflash reporter assay using conditioned media from *Csf1r.iCre; Ctnnb1^fl/fl^* mice BMMɸ shows significantly higher WNT activity (p<0.005) compared to the WT littermates (*Ctnnb1^fl/fl^*) BMMɸ (Figure 3A, B, Figure S1E). Which was further confirmed by ELISA with the detection of higher WNT levels in a CM from *Csf1r.iCre; Ctnnb1^fl/fl^* mice BMMɸ compared to WT littermates BMMɸ (Figure 3C). qPCR analysis demonstrated that the absence of *Ctnnb1* in mice BMMɸ significantly upregulated expression of 14 WNTs in *Csf1r.iCre; Ctnnb1^fl/fl^* mice compared to the *WT* mice (Figure 3D). In this study, for the first time, we showed that the genetic ablation of β-Catenin enhances the WNT expression in macrophages.

**Figure 3:**
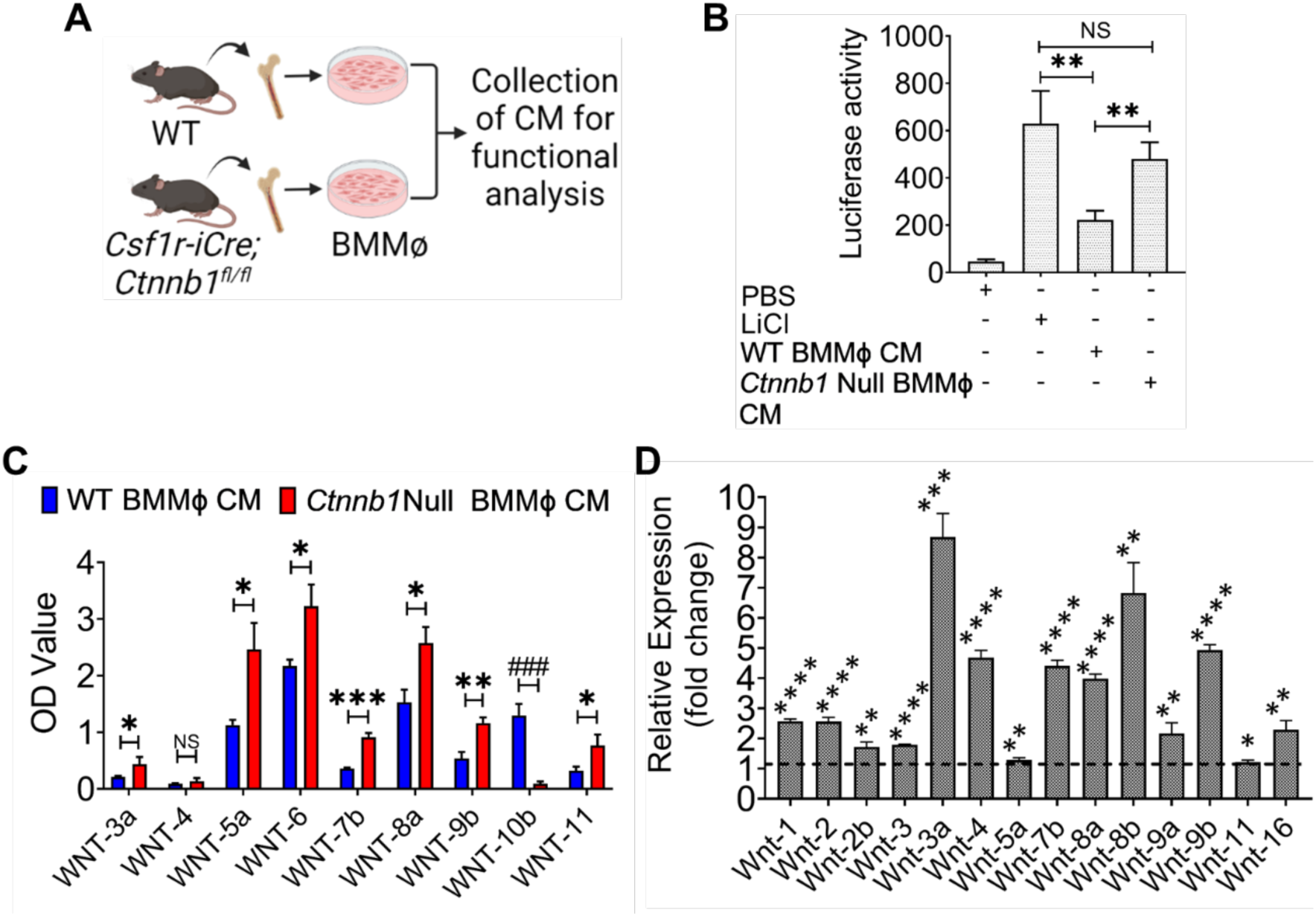
Deletion of gene expressing β-Catenin (Ctnnb1) upregulates expression of macrophage-derived WNT release: (A) Schematic representation of mice BMMɸ culture. (B) TOPflash assay showed increase in WNT activity in BMMɸ CM from Csf1r.iCre; Ctnnb1fl/fl mice compared to WT mice. (C) ELISA showed significance increase in secretory WNT ligands in CM from Csf1r.iCre; Ctnnb1fl/fl BMMɸ compared to WT BMMɸ. (D) qPCR analysis of RNA from Csf1r.iCre; Ctnnb1fl/fl BMMɸ showed significantly higher WNT expression compared to WT BMMɸ. Data presented as the mean ± SD. (Significant level, *: p<0.05, **: p<0.005, ***: p<0.0005, ****: p<0.00005, NS: not significant).

### Modulation of β-Catenin signaling favors the anti-inflammatory phenotype and trophic signals in macrophages

Macrophages are key to innate immunity, tissue homeostasis and regeneration of the damage tissue ^27^. Plasticity of macrophages allows rapid and highly orchestrated responses to various pathogenic and physiological challenges as well in health and disease condition ^28^. To detect the effect of the deletion of β-Catenin gene or pharmacological inhibition of β-Catenin on macrophage phenotype and inflammatory signaling, BMMɸ from *Csf1r.iCre; Ctnnb1^fl/fl^* v/s WT or BMMɸ-XAV v/s BMMɸ were analyzed by RNA-seq. Both pharmacological inhibition and genetic deletion of β-Catenin shifts the balance toward M2 phenotype (Figure S5A-B). Pharmacological inhibition of β-Catenin downregulated 7 genes (Ccl2, Ccl4, Ccl8, Ccl9, Il1b, Tlr4, Socs3) representing M1 macrophage phenotype and upregulated 6 genes (Il4, Tgfb1, Arg1, Cd200, Cd200r1, Ccl22) specific to M2 phenotype. Genetic deletion of β-Catenin downregulated 11 genes (Ccl2, Ccl3, Ccl4, Ccl8, Ccl9, Cd80, Cd86, Tnf, Il1r1, Il1b, Il23r) representing M1 macrophage phenotype and upregulated 6 genes (Il4, Tgfb1, Cd163, Arg1, Cd200, Cd200r1) specific to M2 phenotype. Moreover, macrophages with inhibition or in absence of β-Catenin demonstrated upregulation of anti-inflammatory genes (Cd24a, ler3, Ppard, Rora, Vps35, Gps2, Ldlr, Selenos) and down regulation of inflammatory genes (Fcgr1, Grn, Ctsc, Zbp1, Alox5ap, Stap1, ccl3, Gprc5b, Ptgs2 etc.). In addition, these macrophages showed upregulation of the growth factors (Manf, Cxcl1, Vegfb, Gpi1, Gmfg, Igf1, Tgfb1, Fgf2) making them more efficient for trophic function such as tissue homeostasis and regeneration.

We have also confirmed that overall survival and macrophage function such as phagocytosis was not influenced by pharmacological inhibition of β-Catenin (Figure S6).

### *Csf1r.iCre; Ctnnb1^fl/^*^fl^ mice demonstrated increase in WNT expression in intestinal macrophages and minimize intestinal epithelial radiosensitivity

Tissue macrophages consists of both BM derived and local yolk sac derived macrophages with distinct phenotype and overlapping function ^29^. Our observations analyzing BMMɸ so far suggest that β-Catenin deletion in macrophages induces expression of macrophage derived WNTs. Therefore, it is also important to determine the regulation of WNT expression in tissue macrophages. Our previous study on regenerative role of macrophage derived WNT in intestine demonstrated that both newly recruited and differentiated tissue macrophages express WNT ^2^. According to previous reports macrophages present in pericryptal region and lamina propria delivers critical growth factors and signals including WNT for intestinal epithelial homeostasis ^30,31^ and repair^9,32,33^.

In this study we first determined the WNT expression in total intestinal macrophage population in *Csf1r.iCre; Ctnnb1^fl/fl^* and WT mice. qPCR analysis of flow sorted F480 positive intestinal macrophages demonstrated significant increase in 9 WNTs mRNA expression in *Csf1r.iCre; Ctnnb1^fl/fl^* mice compared to WT littermates (Figure 4A). Though macrophage specific deletion of gene expressing β-Catenin did not show any changes in phenotype of intestinal macrophage population compared to WT littermates (Figure 4B, C).

**Figure 4:**
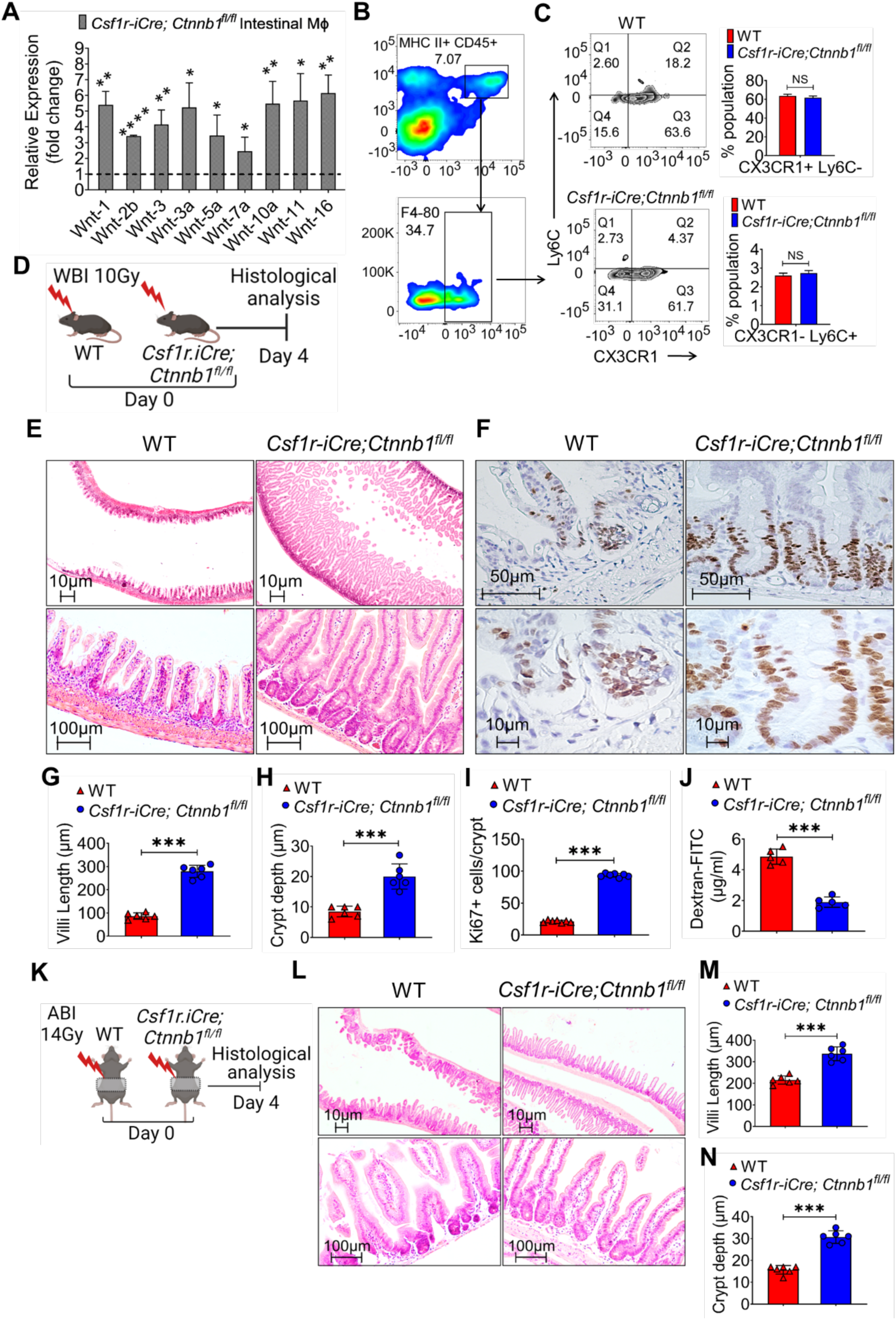
Csf1r.iCre; Ctnnb1^fl/fl^ mice with WNT enriched mucosal macrophages are resistant to RIGS: (A) qPCR of RNA from flow-sorted mice intestinal macrophages. (B-C) Flowcytometry of intestinal macrophages showed no changes in Ly6C+Cx3CR1- and Ly6C-Cx3CR1+ phenotype in Csf1r.iCre; Ctnnb1^fl/fl^ mice vs WT mice. (D) Schematic representation of radiation exposure and timeline for histopathology. (E) Hematoxylin Eosin staining of jejunum section from Csf1r.iCre; Ctnnb1^fl/fl^ and WT mice exposed to 10 Gy WBI (upper panel 2x, lower panel 10x magnification). (G-H) Histogram showing higher villi length (μM) and crypt depth (μM) of jejunal sections from Csf1r.iCre; Ctnnb1fl/fl mice compared to WT. (F) Ki-67 immunohistochemistry in irradiated mice jejunal crypt (upper panel 20x, lower panel 40x magnification). (I) Histogram demonstrating higher number of Ki67+ cells per crypt in Csf1r.iCre; Ctnnb1^fl/fl^ mice compared to WT. (J) Histogram demonstrating low serum dextran level in irradiated Csf1r.iCre; Ctnnb1fl/fl mice compared with WT. (K) Schematic representation of ABI and timeline for histology. (L) Hematoxylin Eosin staining of jejunum section from Csf1r.iCre; Ctnnb1fl/fl and WT mice exposed to 14 Gy ABI (upper panel 2x, lower panel 10x magnification). (M-N) Histogram representing higher Villi length (μM) and crypt depth (μM) in irradiated Csf1r.iCre; Ctnnb1fl/fl mice jejunal sections compared with WT. Data presented as the mean ± SD. (Significant level, ***: p<0.0005, NS: not significant) (n=5 mice per group).

It is important to assess whether genetic ablation of β-Catenin in macrophages had significant consequences on intestinal development and function. *Csf1r.iCre; Ctnnb1^fl/fl^* mice showed adult intestinal length, intestinal morphology including villi length and crypt depth which is very similar to WT littermates (Figure S4A-F).

Next, we determined the radio-sensitivity of intestinal epithelium in *Csf1r.iCre; Ctnnb1^fl/fl^* and WT mice exposed to 10Gy of whole-body irradiation (WBI) (Figure 4D). The histological analysis of jejunum from these mice showed significantly higher villi length (Figure 4E, G) and crypt depth (Figure 4E, H) with a higher number of proliferating cells per crypt as shown by Ki-67 positivity (Figure 4F, I) in *Csf1r.iCre; Ctnnb1^fl/fl^* mice compared to the WT mice.

Since dextran is unable to cross the intestinal epithelial layer unless it is damaged, dextran in the blood is a good indicator of epithelial damage ^2,34,35^. Blood FITC-dextran levels were measured in *Csf1r.iCre; Ctnnb1^fl/fl^* and WT mice at 4 h after gavage with FITC-dextran solution. As demonstrated in Figure 4J, serum dextran level was significantly lower in *Csf1r.iCre; Ctnnb1^fl/fl^* mice in comparison to the WT mice (p<0.0005) suggesting restitution of epithelial integrity compared with WT mice.

In a parallel study, mice were exposed to clinically relevant abdominal irradiation (ABI) (Figure 4K). Exposure to lethal dose of ABI (14Gy) demonstrated significant reduction the villi length (p<0.0005) (Figure 4L, M) and crypt depth (p<0.0005) (Figure 4L, N) in WT mice, while *Csf1r.iCre; Ctnnb1^fl/fl^* mice shows restitution of both villi and crypt. Macrophage derived WNTs activate β-Catenin in the irradiated crypt cells ^2^. Activation and nuclear translocation of β-Catenin drives a gene-expression program that supports stem cell maintenance and proliferation. β-Catenin staining revealed a significant increase in nuclear localization in the intestinal epithelium of *Csf1r.iCre; Ctnnb1^fl/fl^* mice compared to WT mice following ABI exposure (Figure S7). These results suggest that the absence of *Ctnnb1* and thereby induction of macrophage derived WNTs expression made intestinal epithelium more radio resistance in *Csf1r.iCre; Ctnnb1^fl/fl^* mice compared to the WT mice.

### Enriched secretory WNT from *Csf1r.iCre; Ctnnb1^fl/fl^* BMMɸ mitigates radiation induced intestinal injury

To confirm the regenerative potential of *Csf1r.iCre; Ctnnb1^fl/fl^* BMMɸ, derived secretory WNT, CM from these cells were tested in mice model of RIGS. WT mice were exposed to WBI with a dose level (10Gy) which induces RIGS ^6^ and results 100% lethality within 12-14 days post-irradiation (Figure S4G, H). Transfusion of *Csf1r.iCre; Ctnnb1^fl/fl^* BMMɸ CM (150µl of media containing 200µg of protein) at 24 and 48 hours post-WBI rescued these *Ctnnb1^fl/fl^* mice from radiation lethality and improved survival (p<0.005) (80% of total irradiated mice survived). However, mice receiving *Ctnnb1^fl/fl^* (WT) BMMɸ CM demonstrated moderate improvement in survival (only 30% of irradiated mice survived) (Figure S4H). These results clearly suggest that enrichment in macrophage derived WNT should be beneficial to mitigate radiation induced gastrointestinal syndrome.

EVs are one of the major carriers of secretory WNT ligands ^2,36^. Our previous report demonstrated that conditioned media derived EVs consist of functional WNT ligands ^2^. We, therefore, examined the enrichment of WNT ligands in EVs purified from *Csf1r.iCre; Ctnnb1^fl/fl^* mice BMMɸ conditioned media. TOPflash assay demonstrated that the *Csf1r.iCre; Ctnnb1^fl/fl^* mice BMMɸ conditioned media derived EVs show significantly higher WNT activity compared to WT BMMɸ (p<0.0005) (Figure S4I, Figure S1F). WNT ligands are major regenerative paracrine factors for the repair and regeneration of intestinal epithelium ^37^. Previously, our groups and several others have shown that the intestinal organoid model is suitable to study ISC function ^2,38,39^ and examine the efficacy of multiple regenerative signals ^40^ including WNT ^41^. To determine the functional efficacy of EVs enriched in WNT ligands we have used mice jejunal organoids exposed to radiation. Our previous studies ^4^ demonstrated that radiation exposure with a dose level of 6-8 Gy results in measurable levels of radiation toxicity in mouse intestinal organoids. Jejunal organoids exposed to 6 Gy of irradiation were treated with EVs (20μg/500μl of culture media) derived from *Csf1r.iCre; Ctnnb1^fl/fl^* mice BMMɸ CM. On Day 4 and Day 10, post-irradiation organoids were quantified under a microscope for budding organoid vs total organoid ratio. Irradiated organoids treated with *Csf1r.iCre; Ctnnb1^fl/fl^* BMMɸ conditioned media derived EVs (Figure S4K-L) show a significant improvement on day 4 (p<0.05) and on day 10 (p<0.0005) in the budding organoid/total organoid ratio compared to the irradiated control group. We have also determined the effect of irradiation on overall survival of these organoids by using ATP uptake assay ^42^. Irradiation dose of 6 Gy reduces the viability of these intestinal organoids which recovered significantly after treatment with *Csf1r.iCre; Ctnnb1^fl/fl^*BMMɸ conditioned media derived EVs (p<0.0005) (Figure S4M). However, EVs from *Csf1r.iCre; Ctnnb1^fl/fl^* BMMɸ treated with C59 (porcupine inhibitor) ^2^ did not show any WNT activity in TOPFLASH assay and failed to rescue intestinal organoids from radiation toxicity (Figure S4N-O). These results clearly suggest that EVs from *Csf1r.iCre; Ctnnb1^fl/fl^* BMMɸ are enriched with functional WNTs ligand which helps in the rescue of ISCs from radiation toxicity and thereby improves the survival of intestinal organoids.

### Adoptive cell therapy with pre-conditioned mouse bone-marrow macrophage mitigates radiation-induced gastrointestinal syndrome

Adoptive cell therapy has become very popular as regenerative therapy against multiple acute and chronic injury ^43–45^. Both pharmacological and genetic inhibition of β-Catenin induces WNT expression. Considering the potential for therapeutic advancement of these pre-conditioned macrophages we prioritize functional studies with β-Catenin antagonist XAV-939 treated macrophages rather adopting genetic ablation of β-Catenin. To examine whether these pre-conditioned macrophage therapy mitigate RIGS, C57BL/6 mice were exposed to lethal dose of WBI followed by treatment (intravenously) with these macrophages at 24 and 48 hours post-WBI (Figure 5A-C). Irradiated mice without cell therapy showed 100% mortality within 12 days of irradiation with characteristic signs and symptoms of RIGS, including diarrhea (Figure S8A), black stools and weight loss (Figure 5B). Mice receiving XAV-939 treated BMMɸ had well-formed stools, maintained body weight, and had 80% survival beyond 30 days (p<0.005; log rank (Mantel Cox test)). However, treatment with untreated BMMɸ only rescued 40% of mice from radiation lethality (p<0.05 log rank (Mantel Cox test)) (Figure 5C). In consistency to our previous report ^6^ our present study also demonstrated the regenerative response of macrophages against radiation induced lethality. However, present observation suggested that optimum benefit can only be realized once macrophages are pre-conditioned with β-Catenin antagonist.

**Figure 5:**
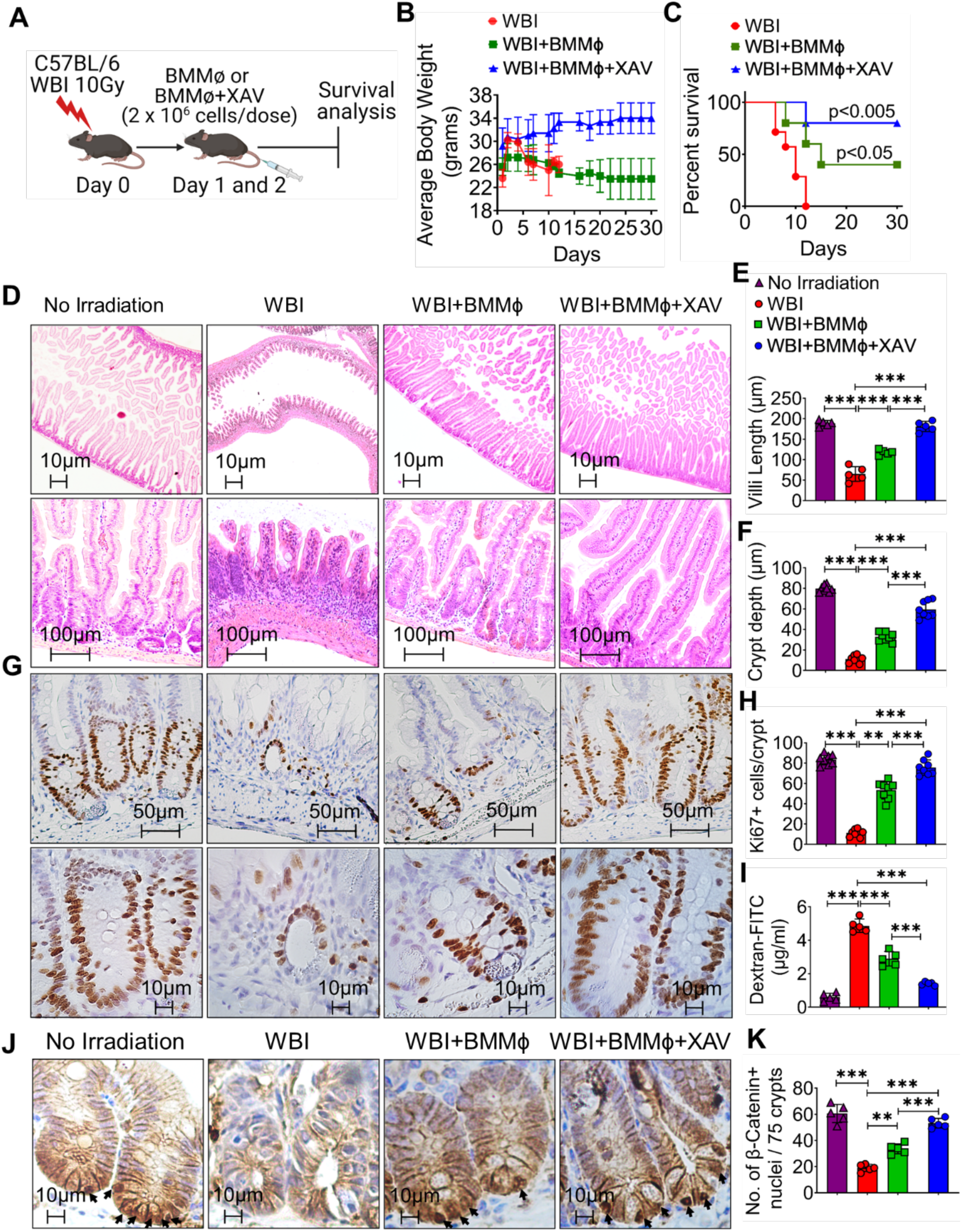
Treatment with pre-conditioned mouse BMMɸ mitigates RIGS: (A) Schematic representation of experimental plan. (B) Body weight of mice at post irradiation time points (n=15 mice per group). (C) Kaplan–Meier survival (Mantel-Cox test) analysis of C57BL/6 mice exposed to WBI (10Gy) shows 80% survival (p<0.005) after pre-conditioned BMMɸ treatment compared to 40% survival (p<0.05) in mice receiving untreated BMMɸ. (D) Hematoxylin and Eosin staining of jejunum section from mice (upper panel 2x, lower panel 10x magnification). (E-F) Histogram showing villus length and crypt depth in jejunal section of C57BL/6 mice. Pre-conditioned BMMɸ treatment showing less reduction of villi length compared to irradiated mice receiving native BMMɸ or no cell therapy. (G) Ki-67 immunostaining of jejunal section (upper panel 2x, lower panel 10x magnification). (H) Histogram demonstrating higher number of Ki67+ cells per intestinal crypt in irradiated mice treated with pre-conditioned BMMɸ. (I) Histogram demonstrating significant low serum dextran level in mice treated with pre-conditioned BMMɸ compared to irradiated group (n=5 mice per group). (J) Representative microscopic images (x40 magnification) of jejunal sections immune-stained with anti β-Catenin antibody to determine β-Catenin nuclear localization. Nucleus stained with hematoxylin. Pre-conditioned BMMɸ treatment showed more nuclear β-Catenin staining (dark brown; indicated with arrows), compared to mice receiving native BMMɸ or no cell therapy. (K) Nuclear β-Catenin count: each data point is the average of the number of β-Catenin-positive nucleus from 15 crypts per field, 5 fields per mice. Number of β-Catenin-positive nucleus in irradiated C57BL/6 mice receiving pre-conditioned BMMɸ is higher compared to mice receiving native BMMɸ or no cell therapy. Treatment with native BMMɸ or without cell therapy following irradiation showed significantly fewer β-Catenin-positive nuclei than the non-irradiated control. Data presented as the mean ± SD. (Significant level, **: p<0.0005, ***: p<0.0005)

The histological analysis of the mice jejunum at 3.5 days post irradiation showed crypt depletion and a decrease in crypt regeneration followed by villi denudation suggesting radiation induced mucosal damage. However, irradiated mice treated with pre-conditioned BMMɸ demonstrated significant restitution of crypt villous structure with an increase in number of villi length (p<0.0005 unpaired *t*-test, two-tailed) (Figure 5D, E) and crypt depth (p<0.0005 unpaired *t*-test, two-tailed) (Figure 5D, F) compared to irradiated control. Irradiated mice receiving untreated BMMɸ failed to show similar level of restitution in mucosal crypt villus structure. Maintenance of mucosal epithelial structure post injury demands strong crypt epithelial regenerative response ^45^. The Ki67 staining on mice jejunum at 3.5 days post-irradiation shows a significantly higher number of proliferating epithelial cells per crypt (Figure 5G, H) with pre-conditioned BMMɸ therapy compared to untreated control mice (p<0.0005 unpaired *t*-test, two-tailed). Irradiated mice receiving untreated BMMɸ also showed improvement in epithelial cell proliferation (p<0.005 unpaired *t*-test, two-tailed) compared to irradiated control, but failed to match the proliferation level achieved with pre-conditioned BMMɸ treatment (Figure 5G, H).

On day-7 post irradiation mice blood FITC-dextran levels were measured at 4 h after gavage. Serum dextran level was significantly lower (Figure 5I) in the irradiated mice receiving pre-conditioned BMMɸ in comparison to the irradiated mice without any cell therapy (p<0.0005 unpaired *t*-test, two-tailed) or receiving untreated BMMɸ (p<0.0005 unpaired *t*-test, two-tailed). Moreover, at day 7 post irradiation small intestinal epithelial morphology and total small intestinal length was completely regained to normal level in pre-conditioned BMMɸ treated group suggesting complete recovery from radiation induced damage (Figure S8B-D).

Immunohistochemical analysis of jejunal sections from C57BL/6 mice showed characteristic nuclear β-Catenin staining with 60.6 ± 6.99 of cells being positive for nuclear β-Catenin (per 75 crypts) (Figure 5J, K). An isotype IgG control is used as a negative control which indicates lack of staining and thus showing specificity for the anti-β-Catenin antibody (Figure S8E). Irradiated C57BL/6 mice had significantly fewer nuclear β-Catenin-positive cells (19 ± 2.74) compared with non-irradiated untreated control (p<0.0005, unpaired t-test, two-tailed), respectively. Treatment with pre-conditioned BMMɸ further improved the number of nuclear β-Catenin-positive crypt epithelial cells (53.2 ± 3.7) compared with irradiated control or irradiated mice receiving untreated BMMɸ (P<0.0005 and P<0.0005, respectively, unpaired t-test, two-tailed) (Figure 5J, K).

### Preconditioned BMMɸ derived EVs mitigates RIGS

Our previous reports demonstrated that intravenous injection of bone marrow derived ^6^ or peripheral blood derived cells ^46^ primarily lodges to lung and few other organs such as bone marrow and intestine and deliver their signal through paracrine manner. We have also shown that mitigation of acute radiation syndrome primarily depends on supplementation of key regenerative signals through paracrine action rather cell replacement therapy to target tissue ^6^. Considering EVs as major carrier of secretory WNT ^47,48^ it is expected that transplanted BMMɸ also deliver the WNT through EV. Therefore, mice receiving pre-conditioned BMMɸ should demonstrate enrichment of WNT in circulating EVs in plasma. EVs purified from mice plasma at 24 hours post cell therapy were tested for the presence of functional WNT. TOPflash assay clearly demonstrated that plasma derived EVs recovered from mice receiving pre-conditioned BMMɸ has significantly higher level of WNT activity compared to mice treated with/without native BMMɸ (Figure S1G).

EV-based therapies are now being prioritized as an alternative to cell transplantation ^49^. Preclinical studies suggested that EV based therapy may overcome the clinical side effects of cell therapy including immune rejection, graft vs host diseases (GVHD) ^50^. In the present study we examined the regenerative potential of BMMɸ derived EV in mice model of radiation induced intestinal injury. According to FDA recommended radiation model for gastro-intestinal acute radiation syndrome C57BL/6 mice was exposed to lethal dose of partial body irradiation (PBI) (12Gy; LD100/14) keeping one hind leg out of the irradiation field for partial bone marrow sparing. Irradiated animals were treated with EVs from pre-conditioned or un-treated BMMɸ CM at 24- and 48-hours post irradiation (Figure 6A). Mice receiving pre-conditioned BMMɸ derived WNT enriched EVs showed significant improvement in survival (80%) (p<0.0005, log rank (Mantel Cox test)) compared to the mice receiving EVs from untreated BMMɸ (50% survival) (p<0.05, log rank (Mantel Cox test)) (Figure 6B). Mice receiving EVs did not show any other clinical side effects and maintained body weight (Figure 6C) suggesting that EV based treatment is safe and can be considered as radiation countermeasure against RIGS.

**Figure 6:**
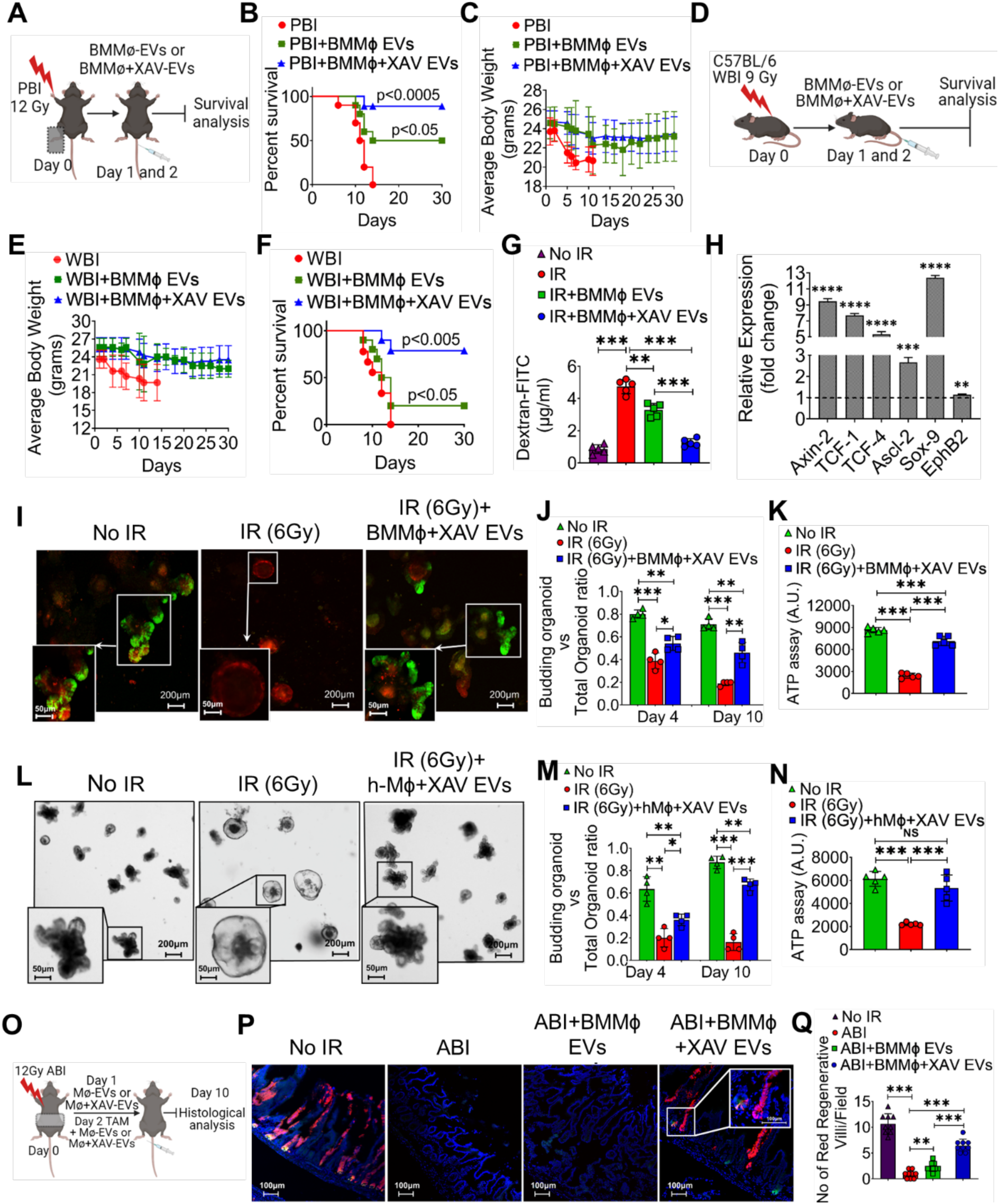
EVs from pre-conditioned BMMɸ conditioned media mitigates RIGS: (A) Schematic representation of PBI and EV treatment. (B) Kaplan–Meier survival (Mantel-Cox test) analysis of C57BL/6 mice receiving pre-conditioned or native BMMɸ derived EVs (200µg) at 24hrs and 48hrs post PBI (12 Gy). Mice receiving pre-conditioned BMMɸ derived EVs showed 80% survival compared to mice receiving native BMMɸ EVs (p<0.05; Log-rank (Mantel–Cox) test) or untreated mice (p<0.0005; Log-rank (Mantel–Cox) test) (n=10 per group). (C) The body weight analysis showed improvement with EVs treatment. (D) Schematic representation of WBI and EV treatment. (E) The body weight analysis showed improvement with EV treatment. (F) Kaplan–Meier survival (Mantel-Cox test) analysis of C57BL/6 mice exposed to WBI (9Gy) shows significant (80%) (p<0.0005) improvement in survival after treatment with pre-conditioned BMMɸ EVs. (n=10 mice per group) compared to mice receiving native BMMɸ EVs (p<0.05; Log-rank (Mantel–Cox) test) or untreated mice (p<0.0005; Log-rank (Mantel–Cox) test) (n=10 per group). (G) Histogram demonstrating significantly low serum dextran level in irradiated mice treated with pre-conditioned BMMɸ EVs compared to irradiated control (n=5 mice per group). (H) qPCR analysis of β-Catenin target genes from C57BL6 mice crypt epithelial cells. (I) Confocal microscopic images of crypt organoids from Lgr5-EGFP-CRE-ERT2; R26-ACTB-tdTomato-EGFP mice. Pre-conditioned BMMɸ EVs rescued the LGR5 +ve (GFP +ve cells) ISCs from radiation injury. Histogram demonstrating the improvement in irradiated organoid growth (J) and ATP uptake (K) with pre-conditioned BMMɸ EVs treatment compared to untreated irradiated control. (L) Microscopic images of organoids from human intestinal surgical specimens. (M) Histogram demonstrating the higher budding organoid ratio in EVs treated group compared to irradiated control. (N) Histogram demonstrating significant increase in ATP uptake with pre-conditioned BMMɸ derived EVs treatment compared to untreated irradiated control. (O) Schematic representation of the treatment plan for lineage tracing assay. (P) Confocal microscopic images of the jejunum section from Lgr5-eGFP-IRES-CreERT2; Rosa26-CAG-tdTomato mice. tdTomato (tdT)-positive cells and Lgr5+ GFP+ cells are shown in red and green respectively. Nuclei are stained with DAPI (blue). (Q) Histogram representing the number of tdT-positive cells containing villi. Pre-conditioned BMMɸ EVs treatment increase in red cell containing villi (n=3 mice per group) compared to native BMMɸ EVs treatment or irradiated control. Data presented as the mean ± SD. (Significant level, *: p<0.05, **: p<0.005, ***: p<0.0005, ****: p<0.0005).

We repeated this study in mice exposed to WBI (Figure 6D). Mice receiving pre-conditioned BMMɸ CM derived EVs at 24- and 48-hours post irradiation-maintained body weight (Figure 6E) and showed significant improvement in survival (80%) compared to the irradiated untreated mice (p<0.005, log rank (Mantel Cox test)) showing 100% mortality within 15 days of irradiation (Figure 6F). Mice receiving untreated BMMɸ CM derived EVs showed only 20% survival rate (p<0.05, log rank (Mantel Cox test).

Serum dextran level was significantly lower in the mice receiving pre-conditioned BMMɸ CM derived EVs in comparison to the mice receiving untreated BMMɸ CM derived EVs (p<0.0005, unpaired *t*-test, two-tailed) or irradiated mice without EV treatment (p<0.0005, unpaired *t*-test, two-tailed) (Figure 6G). This result further confirms the radiomitigating potential of WNT enriched EVs against RIGS.

Supplementation of WNT ligand activates the canonical β-Catenin signaling in crypt epithelial cells ^2,51^. qPCR analysis of WNT target genes in C57BL/6 mice receiving WNT enriched EVs demonstrated significant upregulation of multiple β-Catenin target genes compared to irradiated untreated control (Figure 6H).

### Pre-conditioned BMMɸ derived EVs rescue intestinal stem cells from radiation toxicity

To examine the effect of EVs derived from pre-conditioned BMMɸ conditioned media in repair and regeneration of ISCs we used *in vitro* primary intestinal organoid culture system using intestinal crypts isolated from the *Lgr5/GFP-IRES-Cre-ERT2*; R26-ACTB-tdTomato-EGFPknock-in mice ^4^ which allow visualization of the ISCs. At a radiation dose level of 6 Gy, most of the Lgr5^+ve^ ISCs had disappeared within 48 h post irradiation, resulting in a significant loss of budding crypts with changes in existing crypt morphology indicating inhibition of ISC growth and differentiation in response to radiation exposure. Organoids treated with EVs (20μg/500μl of culture media) derived from pre-conditioned BMMɸ CM demonstrated significant presence of Lgr5+ve green ISCs (Figure 6I) with significant improvement in the budding organoid/total organoid ratio compared to the irradiated control group (day 4 post irradiation p<0.05, unpaired *t*-test, two-tailed; day 10 post irradiation p<0.005, unpaired *t*-test, two-tailed) (Figure 6J). ATP assay also shows improvement in organoid survival with EV treatment (p<0.0005, unpaired *t*-test, two-tailed) as compared to the irradiated control (Figure 6K). In summary, this ex vivo organoid study clearly demonstrated that EV mediated supplementation of WNT at optimum level can rescue Lgr5^+ve^ ISCs and thereby improve survival of intestinal organoids from radiation toxicity.

Mouse intestinal organoids were also tested for expression of other ISC markers representing both +4 position and crypt base columnar cells (CBCs) ^52,53^. Irradiated organoids receiving pre-conditioned BMMɸ CM-derived EVs demonstrated an increase in mRNA levels of ISC specific markers such as OLFM4, LGR5, BMI-1 compared with irradiated untreated organoids (Figure S8G). Irradiated Organoids treated with pre-conditioned EVs demonstrated a several-fold increase in mRNA expression of β-Catenin target genes, indicating activation of Wnt-β-Catenin signaling (Figure S8H).

To examine the effect of pre-conditioned h-Mɸ CM derived EVs on human intestinal epithelial tissue, surgical specimens collected from normal colon at least 10 cm apart from the malignant site were used to develop an ex vivo crypt organoid. At a dose level of 6 Gy all the budding crypts have disappeared in the organoids. However, organoids treated with pre-conditioned h-Mɸ CM-derived EVs post-irradiation had budding crypts with complete restitution of organoid structure (Figure 6L). Quantification of the budding crypt-like structure demonstrated a significant higher number of budding crypts/total crypt ratio in EVs treated group on day 4 (p<0.05, unpaired *t*-test, two-tailed) and day 10 (p<0.0005, unpaired *t*-test, two-tailed) (Figure 6M). EV treatment also helps in the improvement of organoid survival (p<0.0005, unpaired *t*-test, two-tailed) measured by ATP uptake assay (Figure 6N) in comparison to the irradiated untreated group.

In response to EV treatment irradiated human organoids also demonstrated an increase in mRNA levels of ISC-specific markers such as OLFM4, DCLK-1, LGR5 compared with irradiated untreated organoids (Figure S6C). Moreover, these EV treated organoids demonstrated a significant-fold increase in expression of β-Catenin target genes compared to irradiated untreated control, indicating activation of Wnt-β-Catenin signaling (Figure S6D). These results clearly suggest that purified EVs from pre-conditioned BMMɸ or h-Mɸ conditioned media carries functional WNTs ligand which can mitigate radiation toxicity in ISCs and thereby improves the overall survival of human intestinal organoids. Next, we examined the role of pre-conditioned BMMɸ CM derived EVs on ISC survival and regenerative potential by in vivo lineage tracing assay using Lgr5-EGFP-ires-CreERT2-R26-CAG-tdT mice ^4^. In this mouse tamoxifen-mediated activation of cre-recombinase under the Lgr5 promoter expresses tdTomato in epithelial cells derived from Lgr5-positive ISC. Therefore, quantification of these tdTomato (tdT)-positive cells in irradiated epithelium with or without pre-conditioned BMMɸ or BMMɸ CM derived EVs determines the regenerative response of Lgr5-positive ISCs. Mice were exposed to 12 Gy ABI, followed by treatment with preconditioned or untreated BMMɸ derived EVs (Figure 6O). Tamoxifen treatment demonstrated the presence of tdT-positive cells in the crypt epithelium (Figure 6P, Q) in both EV treated groups. However, number of tdT positive cells are much higher in mice receiving preconditioned BMMɸ derived EVs compared to EVs from untreated BMMɸ (p<0.0005). In irradiated untreated mice, tdT-positive cells are absent, suggesting the loss of regenerative capacity of Lgr5-positive ISCs. (Figure 6P, Q). All this evidence clearly demonstrates that pre-conditioned BMMɸ CM derived EVs are superior to promote intestinal epithelial regeneration by inducing the growth and proliferation of ISCs.

### Pre-conditioned BMMɸ derived EVs reduces DSS induced colitis

To investigate the role of BMMɸ derived EVs in response to acute colitis, we chose to use DSS-induced intestinal injury as our model, a commonly used model of inflammatory bowel disease (IBD) ^54^. 8-10 weeks old male C57BL6 mice were treated with 5% DSS (supplemented in drinking water) and followed up for up to 6 days (Figure 7A). The body weight (Figure 7B) was significantly decreased on D4 (p<0.05) until D6 (p<0.05) of DSS treatment. Overall DAI score was significantly high in DSS treated mice compared to control (Figure 7C, D). However, DSS treated mice receiving pre-conditioned BMMɸ derived EVs treatment demonstrated restitution of body weight with significant reduction in DAI score (Figure 7C, D).

**Figure 7.**
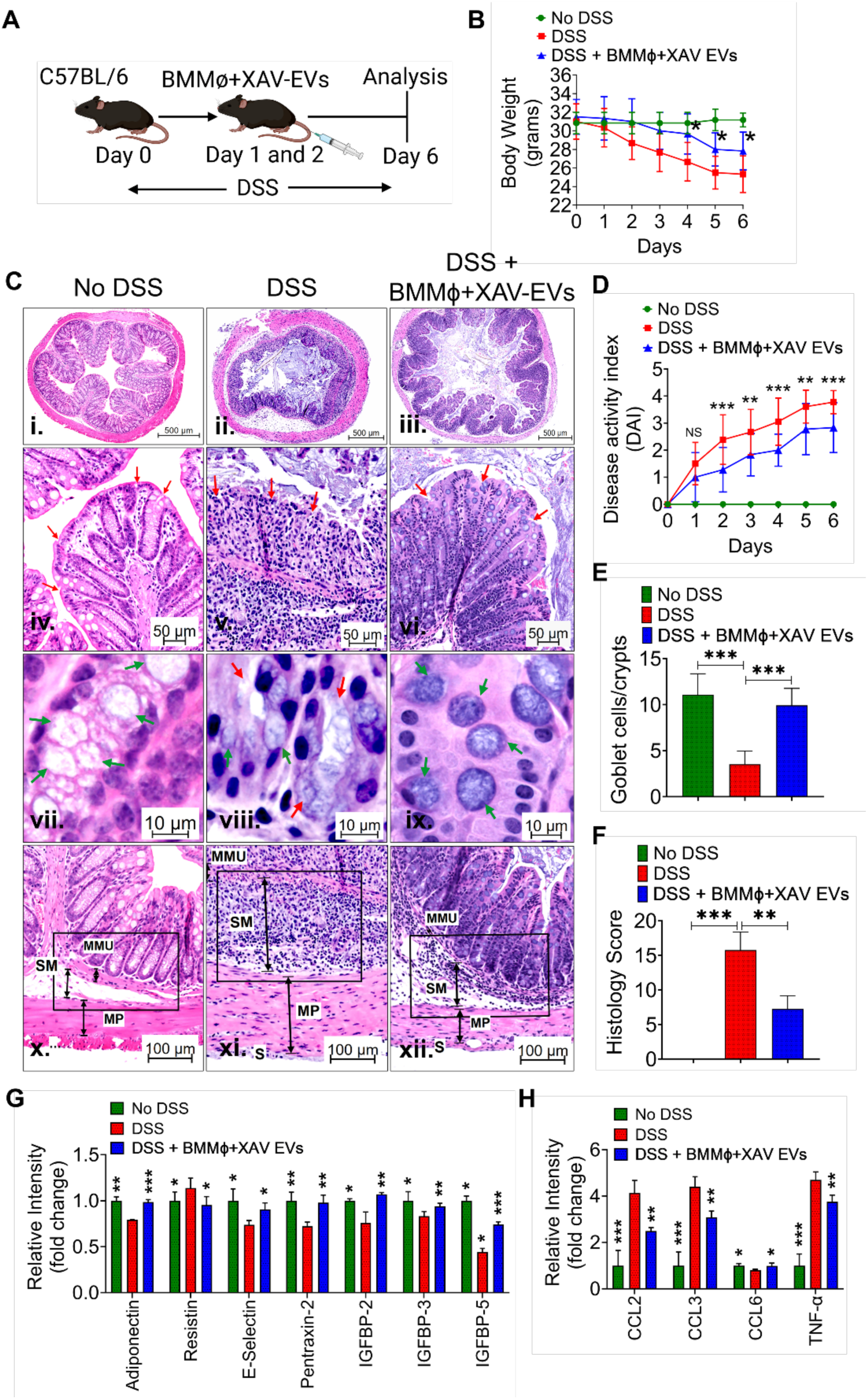
Pre-conditioned BMMɸ derived EVs attenuate DSS-induced colitis in mice. (A) Schematic representation of treatment plan for DSS-induced colitis (B) Changes in body weight over 6 days of DSS treatment, showing recovery with EVs treatment. (C) Representative H&E-stained colonic sections showing mucosal damage (panel iv, v, vi with red arrow), distorted colonic glands (panel viii with red arrow) and large immune cell infiltration (panel xi) in DSS model which was improved with EVs treatment (well-formed mucosal folds (panel vi with red arrow), improved colonic glands with goblet cells (panel ix with green arrow) and very less immune cell infiltration (panel xii). (D) Disease Activity Index scores indicating reduced DSS induced severity with EVs treatment. (E) shows number of mucus secreting goblet cells per crypt (F) Evaluation of histological scores considering goblet cell loss, mucosal damage, crypt density and inflammatory infiltrate reductions after EVs treatment. (F) Cytokine analysis showing improved levels of growth factors and reduced inflammatory cytokines in EVs-treated mice. Data presented as the mean ± SD. (Significant level, *: p<0.05, **: p<0.005, ***: p<0.0005).

Histological analysis with H&E staining showed the severity of colitis was increased from D1 to D6 after DSS treatment (Figure 7C panel ii), the injury included severe colonic epithelium damage with absence of mucosal folds (Figure 7C panel v), distortion in mucus secreting colonic gland structure (shown by red arrow) and loss of goblet cells in gland (green arrow) (Figure 7C, E panel viii). DSS treated mice shows severe immune cells infiltration with extended submucosa (Figure 7C panel xi, black rectangle), and the overall histology scores were highly increased on D6 compared with controls (No DSS treated mice) (p<0.0005) (Figure 7F). However, pre-conditioned BMMɸ derived EVs treatment in DSS treated mice demonstrated less mucosal damage with well-marked mucosal folds (Figure 7C panel vi, red arrow), presence of large number of mucus secreting colonic gland with goblet cells (Figure 7C, panel ix, shown by green arrow). EVs treatment significantly reduces the immune cells infiltration with less submucosal thickness (Figure 7C, panel ix, shown by green arrow) with significant improvement (p<0.005) in overall histology score compared to the DSS treated mice (Figure 7F). The cytokine analysis of the blood plasma collected at the end of study on D6 shows various growth factors/hormonal proteins (Adiponectin, Resistin, E-selectin, Pentraxin-2, IGFBP-2/3/5) was significantly improved after EVs treatment in DSS treated mice (Figure 7G). Furthermore, inflammatory cytokine was also significantly reduced in EVs treated mice (CCL2, CCL3, TNF-α) (Figure 7H). Which suggests that pre-conditioned BMMɸ derived EVs treatment mitigated DSS-induced colitis in mice to some extent, as evidenced by improvements in body weight, DAI scores, histological features/scores, and cytokine profiles. These findings suggest the therapeutic potential of pre-conditioned BMMɸ derived EVs in the treatment of inflammatory bowel diseases.

### Supplementation of WNT ligands does not promote intestinal tumor growth

Enhancement of cancer risk or promotion of tumor growth has always been a concern for any regenerative therapy. Mitigation of radiation toxicity primarily in clinical scenario should be targeted precisely to normal tissue without supporting tumor growth. In the present study we have examined the effect of EV based supplementation of WNT ligand on intestinal tumor growth in ApcMin/+ mice. ApcMin/+ mouse consists a mutation in the Apc gene, responsible for the inherited colon cancer syndrome, familial adenomatous polyposis coli, in humans ^55^. Mouse were treated with pre-conditioned BMMɸ EVs at a concentration of 200μg/dose per mouse. On day 17 post treatment mice were euthanized and intestinal tissue were analyzed for tumor counts. We found that in Apc min+ mice model of intestinal cancer after EVs treatment shows significant regression of the tumor growth compared to the control untreated mice (Figure S9). Although mechanism of tumor regression in response to EV treatment needs to be investigated but this data clearly suggested that pre-conditioned BMMɸ EVs treatment might be safe to use as a radio-modulator to minimize radiation toxicity in intestine without inducing any support to tumor growth or radiation resistance.

## DISCUSSION

WNT signaling plays an important role in both tissue development and repair process. Our previous report, including reports from other groups ^1,2,56^ demonstrated that macrophages are one of the major sources of WNT ligands. Present study indicated for the first time that canonical WNT signaling suppresses the expression of WNT ligand in macrophages. In our study both genetic and pharmacological inhibition of β-Catenin induces the expression of functional WNT ligand in macrophages. Enrichment of WNT ligands expression was also achieved in human PBMC derived macrophages treated with β-Catenin antagonist Tankyrase inhibitor. CHIP-seq analysis clearly demonstrated that β-Catenin is associated with WNT promoters in macrophages. A validation study in HEK293 cells having luciferase reporter system under WNT promoter also demonstrated regulatory role of β-Catenin in WNT promoter activity. β-Catenin does not have a DNA binding domain ^57^ but can interact with DNA either by transcription factor TCF or through other mediators ^57^. In the present study β-Catenin immunoprecipitate demonstrated the presence of TCF. However, further study is needed to validate the role of TCF in β-Catenin mediated regulation of WNT expression.

Macrophage derived WNTs play critical paracrine signals in tissue development and injury repair process ^40,41^. Our present study indicated that enrichment of macrophage derived WNT will be beneficial to ameliorate radiation toxicity in intestine. Deletion of *Ctnnb1,* the gene expressing β-Catenin specific to macrophages demonstrated a significant increase in WNT expression in mucosal macrophages and significant resistance to radiation induced intestinal epithelial toxicity.

Radiation induced Intestinal toxicity is one of the major forms of acute injury which may cause lethality within days post exposure ^58^. So far there is no approved therapy available to mitigate radiation induced gastrointestinal syndrome. Macrophage based regenerative therapy should be very useful to deliver WNT based paracrine signal to injured tissue ^2,6^. However, enrichment of macrophage derived WNT expression will be crucial to deliver optimum level of WNT ligand and promote the efficacy of macrophage-based therapy. Our study showed that transfusion of preconditioned macrophages enriched with WNT ligand can mitigate RIGS in C57BL/6 mice by activating canonical WNT signaling in the intestinal epithelium.

In the current study circulating EVs recovered from mice receiving preconditioned macrophages demonstrated enrichment of WNT ligands and suggesting EVs as major carrier of WNTs released from transfused cells. Treatment with preconditioned BMMɸ derived WNT enriched EVs successfully mitigated RIGS in mice with improvement in regenerative potential of ISCs. Similar response was observed with human preconditioned macrophage derived EV treatment promoting survival and induction of human intestinal organoids with activation of canonical WNT signaling. These results clearly suggest that supplementation of WNT ligand through EV is a feasible alternative approach to mitigate RIGS.

Based on recent reports it appears that macrophage based regenerative therapy is a developing paradigm against tissue injury ^59^. Our study clearly suggests that enrichment of WNT ligands in macrophages will make these cells very effective vehicles to deliver WNT based regenerative signals. However, cell-based therapy has its limitations. It is primarily successful only in syngeneic settings and not possible in allogenic settings due to graft versus host diseases (GVHD) ^60^. EV based treatment suggests a better alternative as it may be delivered in allogenic settings ^50^. Moreover, mitigation of acute injury to an actively self-renewing tissue like intestine demands rapid repair and regeneration. Cell replacement therapy is not feasible to match this need due to the time required for engraftment and repopulation. Our study in both radiation and acute chemical colitis model clearly demonstrated that delivery of paracrine signals through EV should be very effective to stimulate the epithelial repair process. Moreover, EV mediated supplementation of WNT ligand may be safe to use as a radio-modulator to improve therapeutic ratio of abdominal radiation as EV treatment did not promote tumor growth in mice model of Apc min intestinal cancer. Considering the current need of WNT based therapy in multiple pathological conditions, our study on regulation of WNT expression and cell/EV based application will advance the field of regenerative medicine.

## STAR★METHODS

### Animals

To study the role of β-Catenin in macrophage-derived Wnt expression, *Ctnnb1 the* gene expressing β-Catenin was deleted by ablating a floxed allele of *Ctnnb1* with a mononuclear phagocyte-restricted Cre-recombinase expressed under the colony-stimulating factor-1 receptor (*Csf1r*) promoter (*Csf1r.icre*). *Tg (Csf1r.iCre) Jwp.-Ctnnb^fl/fl^* (Csf1r-iCre; Ctnnb1flox) mice were generated by crossing *B6.129-Ctnnb1tm2Kem* (Jackson Labs #004152) with *Tg(Csf1.icre)jwp* (Jackson Labs #021024) ^26^ mice to generate *Csf1r.iCre; Ctnnb1^fl/fl^* mice that will have a deletion of β-Catenin (*Ctnnb1*) gene restricted to macrophages. Thereafter mice were inter-bred and *Cre*-deleted mice compared with littermate *Cre-*negative controls referred to as WT. 8-12 weeks-old male C57BL6/J mice, Lgr5-eGFP-IRESCreERT2 mice, Gt (ROSA)26Sortm4(ACTB-tdTomato-EGFP) Luo/J mice, and B6. Cg-Gt (ROSA)26Sortm9(CAGtdTomato)Hze/J mice were purchased (Jackson laboratories) for experimental studies. Lgr5-eGFP-IRES-CreERT2 mice were crossed with Gt (ROSA)26Sortm4(ACTB-tdTomato-EGFP) Luo/J mice (Jackson Laboratories) to generate Lgr5/eGFP-IRES-Cre-ERT2; R26-ACTB-tdTomato-EGFP ^4,61^ and In Gt (ROSA)26Sortm4(ACTB-tdTomato-EGFP) Luo/J mice tdTomato is constitutively expressed (independent of Cre recombination) in the membrane of all cells, and therefore allows better visualization of cellular morphology. Lgr5-eGFP-IRES-CreERT2 mice were also crossed with B6. Cg-Gt (ROSA)26Sortm9(CAG-tdTomato) Hze/J mice (Jackson Laboratories) to generate the Lgr5-eGFP-IRES-CreERT2; Rosa26-CAG-tdTomato heterozygote for lineage tracing. All the animals were maintained ad libitum, and all studies were performed under the guidelines and protocols of the Institutional Animal Care and Use Committee of the University of Kansas Medical Center. Animal studies were performed under the experimental protocols approved by the Institutional Animal Care and Use Committee of the University of Kansas Medical Center (ACUP number 2019-2487).

### Isolation, culture, and pre-conditioning of Mouse Bone marrow derived, and Human Peripheral Blood Mononuclear Cells derived Macrophages

BM was isolated from both femurs and tibias of adult females *Csf1r.iCre; Ctnnb1^fl/fl^* mice and their WT littermate control as well as from the C57BL/6 mice ^62^. The bones were flushed with BMMɸ culture media containing αMEM (Gibco) with 10% (v/v) fetal bovine serum (FBS), 1% penicillin-streptomycin, and 1% Glutamax. The isolated BM cells were plated into 100 cm^2^ tissue culture plates with culture media supplemented with 100ng/ml of M-CSF (Cat no. 315-02-250, Peprotech Inc., NJ, USA) for 24 h. To generate fully differentiated monocyte-derived macrophages non-adherent cells were transferred to petri dishes and cultured for 6–7 days at 37^0^C with 5% CO2 with culture media supplemented with 100ng/ml of M-CSF.

For preconditioning BMMɸ were treated with/without various concentration of XAV-939 (10-20μM) in complete expansion media with incubation at 37^0^C for two hours. Thereafter cells were washed three times with PBS to remove all the traces of XAV-939. Untreated or treated cells were either harvested for detection of β-Catenin by western blot analysis or cell therapy study (2x 10^6^ cells/ dose/ mice, total 2 doses/ mice) or continued in culture for collection of CM subjected to TOPflash or in vitro organoid assay. For TOPflash assay BMMɸ culture was continued for 24 hours in presence of complete alpha MEM (without phenol red) prior to CM collection. For organoid assay BMMɸ culture was continued for 48 h in serum deprived medium (0.5% v/v FBS) before collection of CM. Prior to experimental use CM was collected and concentrated (10-fold) with centrifugal filter units (Millipore, Billerica, MA) ^63,64^.

h-Mɸ were cultured from PBMCs purchased from StemCell Technologies (Cat no. 70025.3). PBMCs were plated in T75 flasks. After 24 hrs of incubation, cells were transferred to a fresh tissue-culture flask and cultured for seven days with expansion media containing StemSpan™-XF with addition of Expansion Supplement (StemCell Technology, Vancouver, BC, Canada) and 100ng/ml of h-CSF (Cat No. 574808, BioLegend, San Diego, CA, USA). Following maturation h-Mɸ were treated with/without XAV-939. After treatment with XAV-939 h-Mɸ were continued in culture to collect CM according to protocol described for BMMɸ.

### Purification of EVs from Mɸ Conditioned media

EVs were isolated from the BMMɸ or h-Mɸ CM adhering to the published guideline ^63,64^ using ultracentrifugation method ^46^. Briefly, CM was centrifuged at 500 x g for 10 minutes at 4^0^C to remove the cells. The supernatant was centrifuged in 2000 x g for 10 minutes at 4^0^C to remove any leftover dead cells. To remove the apoptotic bodies/debris, the supernatant was centrifuged at 10,000 x g for 30 min at 4^0^C. The supernatant was then subjected to ultracentrifugation at 100,000 x g for 70 min at 4^0^C. The pellet containing EVs was washed with ice-cold PBS at 100,000 x g for 70 min at 4^0^C. The EVs pellet was resuspended in PBS and stored at −80^0^C for further studies. The amount of EV protein recovered was assessed using the micro bicinchoninic acid (BCA) assay kit (Thermo Scientific, Waltham, MA, USA) according to the manufacturer’s instructions. The particle count, size distribution and concentration of isolated EV were measured by Nanosight NTA (LM10, Malvern Inst. Ltd., UK). The NTA analyzes the motion of particles illuminated by a laser, from which it deduces their size and concentration. Purified EVs at the concentration of 100 μg/ml, 200 μg/ml and 200 μg/dose/mice (total 2 doses/mice) were used for TOPFLASH assay, ex vivo organoid assay and in vivo treatment respectively.

### Characterization of EVs

Mouse and human macrophage CM derived EVs were characterized by flowcytometry ^63,64^ using mouse and human macrophage specific FITC CD9 (Clone H19a, BioLegend), Pe CD81 (clone 5A6, BioLegend), APC CD63 (clone H5C6, BioLegend) antibody cocktail in 30 µL of suspension and incubated 20 min at 4^0^C in the dark. After incubation, the EVs were washed twice with PBS and used for flowcytometric analysis. Flowcytometric data were analyzed by FlowJo software.

### Irradiation procedure

Whole-body irradiation (WBI) was performed on mice anesthetized with 87.5 mg/kg of Ketamine and 12.5 mg/kg of Xylazine using a small animal radiation research platform (XENX; XStrahl) (0.67 mm Cu HVL) at a dose rate of 3.1 Gy/min at 220 kVp and 13 mA as previously described ^46^. For abdominal irradiation (ABI), a 3-cm^2^ area of the mouse abdomen encompassing the GI tract was irradiated at a dose rate of 2.26 Gy/min at 220 kVp and 13 mA, but shielding the upper thorax, head, and neck as well as the lower and upper extremities ^4^. This results in significant protection of the bone marrow, while predominantly inducing RIGS. For partial body irradiation (PBI), the mouse was irradiated at a dose rate of 2.82 Gy/min at 220 kVp and 13 mA, by shielding one leg from radiation exposure ^65^. To ensure homogeneous dose delivery, half of the dose was delivered from the anterior-posterior direction and the other half from the posterior-anterior direction. The total irradiation time to deliver the intended dose was calculated with respect to dose rate, field size for radiation, and fractional depth dose to ensure accurate radiation dosimetry.

### Mouse colitis model

Colitis was induced in C57BL/6 mice by addition of 5% Dextran Sulfate sodium (DSS) colitis grade (weight/vol) (Cat no. DS1004, Gojira Fine Chemicals, OH, USA) to the drinking water. Mice were treated with pre-conditioned BMMɸ EVs post 24 and 48 hours of DSS initiation. The body weights of all the mice were assessed every day during the treatment period. At Day-6 histological assessment of colitis was performed by H&E staining while blood plasma was analyzed for cytokine analysis. Disease activity index (DAI) and Histology score was calculated as per the parameters mentioned in table S1.

### TCF/LEF (TOPflash) reporter assay

The HEK293 cells (Signosis, Santa Clara, CA) having TCF/LEF luciferase reporter construct ^4,21^ were treated with macrophage CM or EVs to determine the canonical WNT activity. LiCl (10 mM) treatment was used as a positive control for luciferase activity. After 24hr of treatment luciferase activity was determined using the Bright-Glo™ Luciferase Assay System (Cat. No. E2610, Promega). HEK293 cells having FOPflash construct were used as a negative control having mutated TCF/LEF-binding site. CM prepared in different batches under the same conditions shown consistent WNT activity.

### Chromatin Immunoprecipitation Assay

The cultured C57BL/6 BMMɸ were treated with LiCl (10mM, 4hrs) followed by Chromatin Immunoprecipitation Assay (ChIP assay) using anti-β-Catenin antibody (Cell Signaling Technology, United States) according to the manufacturer’s protocol described in the ChIP assay kit (Cell Signaling Technology, United States). Rabbit IgG (Cell Signaling Technology, United States) antibody was used as a negative control. In brief, Cells were fixed with 1% formaldehyde for 10 min at room temperature and quenched with 125 mM glycine for 5 min at room temperature. The DNA was digested with 0.5 μl of micrococcal nuclease into an average fragment size of 100–900 bp. The sample was incubated overnight at 4^0^C with anti-β-Catenin (Cell Signaling Technology, United States), or the normal rabbit IgG (Cell Signaling Technology, United States) as a control followed by incubation with magnetic beads for 2 hours at 4^0^C with gentle rotation. The immunoprecipitate was collected by placing the tubes in a magnetic rack followed by washing and elution was performed using elution buffer. Further, the cross-links were reversed by incubating the eluted samples at 65^0^C for 2 hours in 5M NaCl and Proteinase K solution. The DNA purification was performed using a spin column. ChIP DNA was processed for DNA sequencing at BGI (BGI Americas Corporation, Cambridge, MA, USA) using BGISEQ-500 platform. Sequencing data was filtered to exclude the low quality reads and clean data reads were mapped to reference genome using SOAPaligner/soap2^66^. After alignment, the genome mapping ratio for the sample was 8,350,329/14,869,364 with mapped rate 56.16%. The quality test was also performed on clean reads number 14,869,364 which represents clean reads Q20 rate 95.42%. This alignment result was used to get the peak length and depth distribution. Later, the peak information was used for bioinformatics analysis. To identify the Wnt promoter sequence only those peak beds were picked up who have overlapped with upstream 2k bp for Wnt genes. Further the identified Wnt promoters were tabulated based on their confidence score (Fig. 2B, Table S2).

To validate the inhibitory role of β-Catenin on Wnt promoter activity we developed a recombinant DNA construct expressing Luciferase reporter under WNT5a and WNT9b promoter. HEK293T cells were transfected with either Scramble plasmid (pTA-Luc, LR-2200, Signosis Inc, CA, USA), or pTA-WNT5a promoter, or pTA-WNT9b promoter (Fig. 2C) (Signosis Inc, CA USA). After 24 hours of post transfection cell-culture media containing DMEM with 10% FBS was added to the wells containing transfected cells and incubated for 24 hours. After 48 hours of post-transfection, samples were processed for CHIP with anti-β-Catenin antibody using the ChIP assay kit (Cell Signaling Technology, United States). The DNA samples obtained from ChIP were subjected to qPCR to determine the presence of WNT5a and WNT9b promoter (Table S2). Quantitative analysis of ChIP qPCR data was presented as fold enrichment over the negative control.

### Development of in-vitro culture of intestinal crypt organoids

Jejunal tissue from Lgr5/eEGFP-IRES-Cre-ERT2; R26-ACTB-tdTomato-EGFP mice, and non-malignant colon tissue from human surgical specimens were used for Crypt cells isolation and development of ex vivo organoid culture as described previously ^2,67^. Human colon tissues were received from the University of Kansas Medical Center Biospecimen Repository (HUS#5929). Mouse jejunum and human colon tissue were scraped and chopped into approximately 5 mm pieces. The tissue was then washed with cold PBS and incubated in a dissociation buffer (Cat No. 100-0485, Stem cell technologies) for 20-30 minutes with agitation at room temperature. For mouse tissue, dissociation buffer was discarded, and fragments were suspended vigorously with a 10-mL pipette in cold 0.1% BSA in PBS, yielding supernatants enriched in epithelial cells. Samples were passed through 70-μm filters (BD Biosciences) to obtain fraction 1. The same step was repeated three times to get fractions 2, 3, and 4. Fractions 3 and 4, enriched in crypt stem cells, were centrifuged at 300g for 5 minutes at 4^0^C and diluted with PBS containing 0.1% BSA for mouse crypt cells.

For human tissue, dissociation buffer was discarded by aspiration after centrifugation at 290g for 5 minutes at 4^0^C. 1ml of ice cold DMEM with 1% BSA was added and vigorously pipetted up and down about 20 times to remove crypts from tissue. Samples were passed through a 70-μm filter into a new tube to get isolated crypts from human tissue.

Crypts were embedded in a Matrigel (Cat No. 356231, Corning) and plated in a prewarmed 24-well plate. After the matrix/Matrigel solidified, mouse and human intestinal organoid Medium (Stemcell Technologies) with their respective supplements and gentamycin (50 mg/mL) were overlaid in their respective wells. The organoid was passage once per week. The number of organoids per well was counted by EVOS Microscope (Thermo Fisher Scientific). Images of organoids were acquired using the fluorescent (Nikon, 80i) and confocal (Nikon, A1RMP) microscopy.

### Histology

The radiation doses more than 8Gy induces cell cycle arrest and apoptosis of the crypt epithelial cells within a day after irradiation which results in a decrease in regenerating crypts in the intestinal tissue by day 3.5 and ultimately villi denudation by day 7 post-radiation exposure ^6^. We sacrificed those irradiated animals when moribund or at 3.5 days after WBI or ABI or PBI and intestines were collected for histopathological analysis. The intestine of each animal was dissected, washed in PBS to remove intestinal contents, and the jejunum was fixed in 10% neutral-buffered formalin prior to paraffin embedding. Tissue was routinely processed, and 5-μm sections were prepared for hematoxylin and eosin (H&E) and IHC staining. All H&E staining was performed at the Pathology Core Facility at the University of Kansas Cancer Center (Kansas City, KS). Details of histopathological analysis include assessment of crypt proliferation rate, villi length and crypt depth.

### Crypt proliferation rate

To visualize the villous cell proliferation, mid-jejunum was collected for paraffin embedding and Ki67 immunohistochemistry. Tissue sections were routinely deparaffinized and rehydrated through graded alcohols and incubated overnight at room temperature with a monoclonal anti-Ki67 antibody (M7240 mib1; Dako). Nuclear staining was visualized using streptavidin-peroxidase and diaminobenzidine (DAB) and samples were lightly counterstained with hematoxylin. Murine crypts were identified histologically according to the criteria as described by Potten at el ^68^. The proliferation rate was calculated as the percentage of Ki67-positive cells over the total number of cells in each crypt. A total of 50 crypts were examined per animal.

### Determination of villi length and crypt depth

Digital photographs from H&E-stained slides were captured in a blind manner using Nikon NIS element 80i microscope. The length from the crypt-villus junction to the villous tip was measured to determine the villous length. Crypt depth and villi length were measured using NIS element 80i microscope software. A total of 15 crypts were examined and 5-10 fields per animal for all histological parameters.

### FITC-dextran permeability assay

Mice exposed to WBI and/or pretreated with BMMɸ or BMMɸ+XAV were gavaged on day 7 after irradiation with FITC-dextran solution (4,000 kDa size, Sigma-Aldrich) (0.6 mg/kg body weight). Mice were euthanized and blood/serum was obtained with cardiac puncture at 4 hours after gavage ^37^. Samples were measured in a 96-well plate using a multiwell fluorometer. A standard curve was constructed using mouse serum having increasing amounts of FITC-dextran to determine the serum levels of FITC-dextran in different treatment groups.

### In vivo lineage tracing assay

To examine the contribution of Lgr5 ISC to tissue regeneration under steady-state conditions, lineage tracing was induced by tamoxifen administration in Cre reporter mice (Lgr5-eGFP-IRES-CreERT2; Rosa26-CAG-tdTomato heterozygote) to mark the ISC and their respective tdT-positive progeny ^41,69–71^. Adult mice were injected with tamoxifen (Sigma; 9 mg per 40 g body weight, intraperitoneally) to label Lgr5+ lineages ^69^. For irradiation injury studies, mice were exposed to 12 Gy ABI, and tissue was harvested on day 10 post-irradiation.

### Isolation of Intestinal Epithelial Cells

To isolate intestinal epithelial cells, Intestinal segment was kept into a 10 cm dish containing 5 mL ice-cold PBS (Ca++ and Mg++ free). Any membrane, blood vessels, and/or fat were removed from the exterior of the intestine. Intestinal content was gently flushed with 1 mL ice-cold PBS by inserting a 1 mL pipette tip into one of the open ends of the intestine segment. The intestinal section was opened lengthwise using small scissors and forceps, intestinal segment was moved through the clean PBS to rinse thoroughly. Then, segment was cut into 2-5 mm pieces, allowing these pieces to fall into the 15 ml 1x PBS in the 50 ml tube. The pieces were settled by gravity and then gently the supernatant was aspirated off, leaving enough liquid to just cover the pieces of tissue. This process was repeated using fresh 15 ml PBS for total 15-20 times or until the supernatant is transparent. The tissue pieces were finally resuspended in 25 mL room temperature Gentle Cell Dissociation Reagent (Stem cell technologies) and incubated at room temperature (15 - 25°C) for 15 minutes on a rocking platform at 20 rpm. Finally, the supernatant was discarded once the tissue segments were settled by gravity at the bottom. The tissue pieces were resuspended in 10 mL ice-cold PBS containing 0.1% BSA and pipette up and down three times. Once the majority of the intestinal pieces are settled to the bottom, the supernatant was passed through 70μm filter and collected in a fresh 50 mL conical tube. This was repeated another 5 times to collect all epithelial cells from top of the villi to crypt. Finally collected supernatants were centrifuged at 290xg for 10 minutes at 2-8°C. The number of viable cells was quantified using an automated Cell Counter T20 (Bio-Rad) with the Trypan blue exclusion method. The isolated cell pellet was used for further analysis.

### β-catenin immunohistochemistry of mouse jejunum

β-catenin immunohistochemistry was performed in paraffin-embedded sections of mouse jejunum. Before immunostaining antigen retrieval was performed by heating slides in pH 6.0 citrate buffer at sub-boiling temperature for 10 minutes in an 1x antigen retrieval solution (Dako, Denmark). Non-specific antibody binding was blocked for 30 min by incubation with 5% blocking buffer (Cat no. 50062Z, Thermofisher) in PBS. Tissue was stained using the anti-β-catenin Antibody (1:100 dilution; Cat no. 610154, BD Transduction Laboratories, Franklin Lakes, NJ;) at room temperature for 2 h followed by staining with horseradish peroxidase-conjugated Anti-Mouse Antibody (Dako, Denmark) at room temperature for 1 h. Peroxidase activity was detected by adding DAB substrate. The nucleus was counter-stained with hematoxylin (blue). β-Catenin-positive nucleus (stained dark brown) was calculated from 75 crypts per animal.

### Phagocytosis Assay

A phagocytosis assay was conducted to investigate the effect of pharmacological degradation of β-catenin using XAV-939 on the phagocytic activity of pre-conditioned human macrophages. The experiment involved the use of two types of conjugate particles: pHrodo™ Green E. coli BioParticles™, and positive control particles pHrodo™ Green Zymosan A BioParticles™, macrophages treated with PBS was used as a negative control. Human macrophages treated with or without XAV-939 were incubated with their respective conjugated particles at a concentration of 0.1 mg/ml for 2 hours at 37°C to allow the particles to interact with the macrophages. After the incubation period, the cells were washed once with PBS to remove any unbound or non-phagocytosed particles. The nuclei of the macrophages were labeled using DAPI mounting media. Confocal microscopy was employed to capture images to assess the phagocytic activity of the XAV-939 treated or untreated macrophages.

### Quantitative Real time PCR

qPCR was performed to determine the WNT ligands expression in mouse and human Mɸ untreated and pre-treated with XAV-939 and Intestinal sorted macrophages from WT and Csf1r.iCre; Ctnnb1fl/fl mice. The presence of WNT promoters in CHIP seq DNA samples. β-Catenin target gene analysis in Intestinal epithelial cells prepared from the jejunum of ABI irradiated of adult male C57BL/6 mice untreated/pretreated with BMMɸ-XAV-EVs. and β-Catenin target genes and stem cells marker expression in mouse and human irradiated intestinal organoid cells treated with respective pre-conditioned Mɸ EVs compared to the irradiated untreated control. Total RNA was isolated from all the samples using RNeasy micro/mini kit from (Qiagen, Germantown, MD, USA). The concentration and purity of the extracted RNA were checked using a NanoDrop spectrometer (Thermo Scientific, Waltham, MA, USA). 1 µg of total RNA was reverse transcribed using RNA to cDNA EcoDry™ Premix (Double Primed) (Takara Bio USA Inc., San Jose, CA, USA). The reaction mixture was incubated for 1 h at 42°C; incubation was stopped at 70°C for 10 min. Quantitative real-time PCR (qPCR) was performed using the QuantStudio™ 7 Flex Real Time PCR System (Applied Biosystems™, New York, NY, USA) and SYBR Green Supermix (Bio-Rad, Hercules, CA, USA) with specific primers to the target genes in a 20 µL final reaction volume. β-Actin/GAPDH was used as a reference gene for sample normalization. The delta-delta threshold cycle (ΔΔCt) method was used to calculate the fold change expression in mRNA level in the samples. All the primer sequences used in this study are listed in Supplementary table S3 are listed here:

### Immunoblot analysis

Mouse or human Mɸ untreated and treated with XAV-939 were analyzed for β-Catenin expression, while mouse BMMɸ following CHIP were analyzed for β-Catenin and/or TCF-4 expression using western blot. For immunoblot analysis, cells were lysed with 1 × RIPA buffer (Cell Signaling, MA, USA) containing a protease and phosphatase inhibitor cocktail (Thermo Fisher Scientific Inc.) to isolate total protein. The cell lysate was centrifuged at 16,000 g for 10 min, and the supernatant was collected in separate tubes. The total protein concentration of the samples was determined by the BCA method. For immunoblot analysis, 60 µg of protein samples were incubated with 1 × gel loading buffer, separated by SDS-PAGE (4–15%, Bio-Rad), and transferred to PVDF membranes using a semi-dry power blotted XL blot transfer system (Invitrogen). After protein transfer, the membrane was blocked with 5% skim milk for one hour followed by incubation overnight at 40C with primary antibody against β-Catenin (1:1000 dilution) (Cell-Signaling, Cat no. 9582) and TCF-4 (Cell Signaling, Cat no:2569T). After incubation with primary and secondary antibodies, the membrane was developed with Trident Femto Western HRP substrate (GeneTex, Irvine, CA, USA) and imaged using a ChemiDoc XRS + molecular imager (Bio-Rad). β-Actin was used as an internal control for normalization.

### RNA Seq Analysis

RNA-Sequencing was performed at a strand specific 100 cycle paired-end resolution, in an illumina NovaSeq 6000 sequencing machine (Illumina, San Diego, CA). The analysis was performed in biological duplicates of the three treatment groups. The 6 samples were multiplexed in an S1 flow-cell, generating between 52.7 and 56.4 million reads per sample. The read quality was assessed using the FastQC software ^72^. On average, the per sequence quality score measured in the Phred quality scale was above 30 for all the samples. The reads were mapped to the mouse genome (GRCm38) using the STAR software, version 2.6.1c ^73^. On average, 92% of the sequenced reads mapped to the genome, resulting between 46.9 and 53.0 million mapped reads per sample, of which on average 84% were uniquely mapped reads. Transcript abundance estimates were calculated using the feature Counts software ^74^. Expression normalization and differential gene expression calculations were performed in DESeq2 software ^75^ to identify statistically significant differentially expressed genes. The significance p-vales were adjusted for multiple hypotheses testing by the Benjamini and Hochberg method ^76^ establishing a false discovery rate (FDR) for each gene.

### Statistics

Mice survival/mortality in different treatment groups was analyzed by Kaplan–Meier (Mantel-Cox) statistics as a function of radiation dose using GraphPad Prism-9.0 software. Mice were sorted randomly after genotyping to each experimental and control group. The minimum number of mice used for survival/mortality study was n=10 per group.

For intestinal sampling regions were chosen at random for digital acquisition for quantitation. Digital image data was evaluated in a blinded manner as to treatment. A two-sided Student’s t-test was used to determine significant differences between experimental groups. All data are presented as mean ± standard deviation (SD). A difference between groups with ∗p<0.05, ∗∗p<0.005, or ∗∗∗p<0.0005 was considered statistically significant.

## ACKNOWLEDGMENT

This work was funded by the NIH/NIAID U01AI138323 to S.S.

## AUTHOR CONTRIBUTIONS

Conceived and design the experiments: S.S., Performed the experiments: R.M.C., P.B., Analysed the data: R.M.C., P.B., R. Z., Radiation dosimetry and Calibration: R.B., Wrote the Paper: R.M.C., P.B., S.S., Edited the manuscript: R.M.C., S.S., Contributed reagents/materials/funding: S.S. All the authors read and approved the final manuscript.

## DECLERATION OF INTERESTS

All authors have no conflict of interest.

## DATA AND MATERIALS AVAILABILITY

All data associated with this study are available in the main text or the Supplementary Materials.

