## supplement file for "Modulation of β-Catenin is important to promote WNT expression in macrophages and mitigate intestinal injury"

**List of Supplementary Materials**

Figures S1 to S9

Tables S1-S3

**Supplementary Figure 1:**

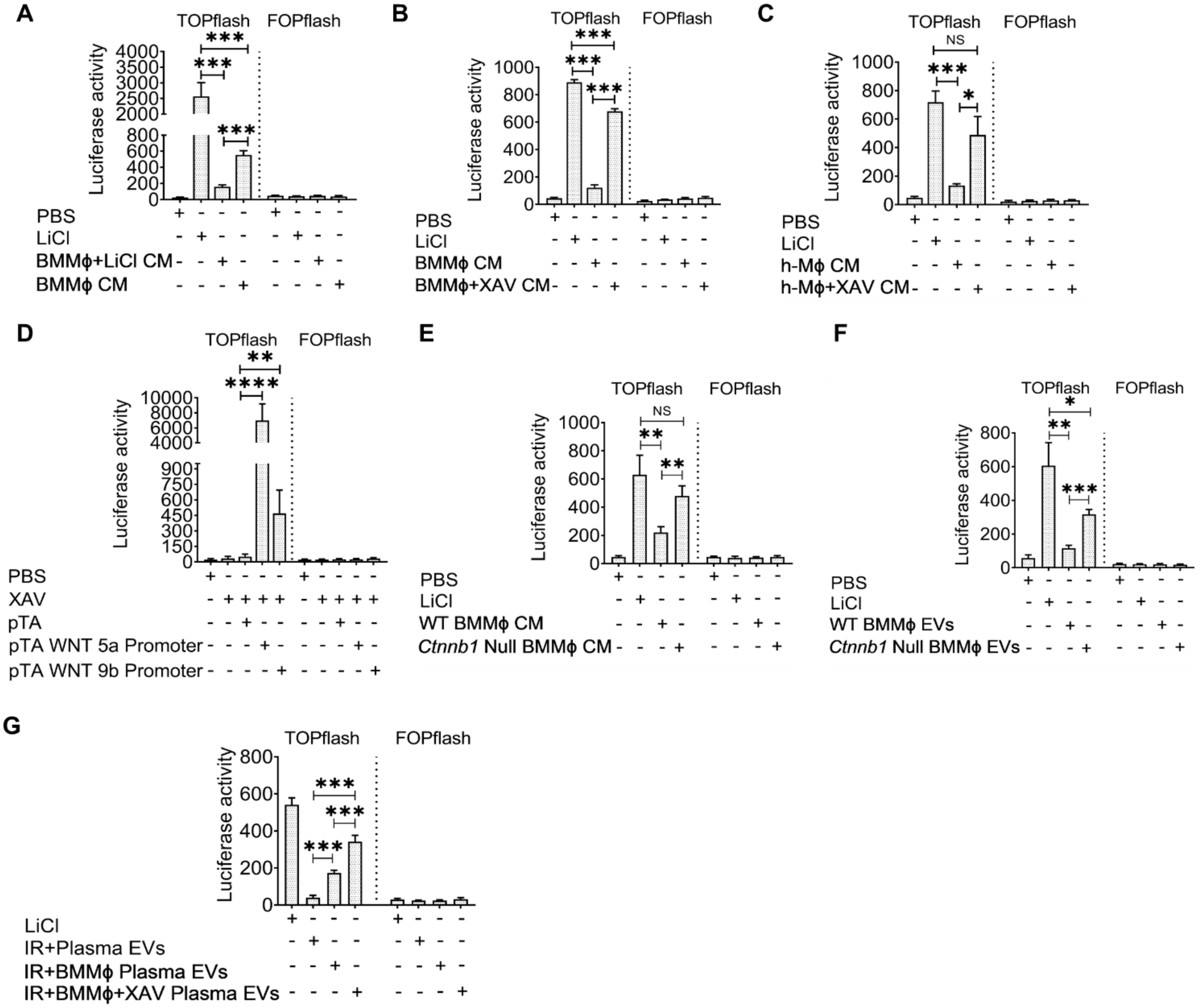


**Supplementary Figure 1: FOPflash luciferase reporter assay:** HEK293 cells having FOPfLASH construct (mutated TCF/LEF-binding site) was used as a negative control for TOPflash assay. Even if *β-Catenin* translocated to the nucleus to activate TCF-mediated transcription, no luciferase activity was seen in the control FOPFLASH reporter assay. (A) FOPFLASH assay of Fig. 1A, (B) FOPFLASH assay of Fig. 1C, (C) FOPFLASH assay of Fig. 1F, (D) FOPFLASH assay of Fig. 2D, (E) FOPFLASH assay of Fig. 3B, (F) FOPFLASH assay of Fig. 3E, (G) TOPflash and FOPflash assay demonstrated enrichment of WNT in circulating EVs from plasma isolated from mice receiving pre-conditioned BMMϕ compared to mice treated with/without native BMMϕ. Data presented as the mean ± SD. (Significant level, *: $p<0.05$, **: $p<0.005$, ***: $p<0.0005$, ****: $p<0.00005$).

**Supplementary Figure 2:**

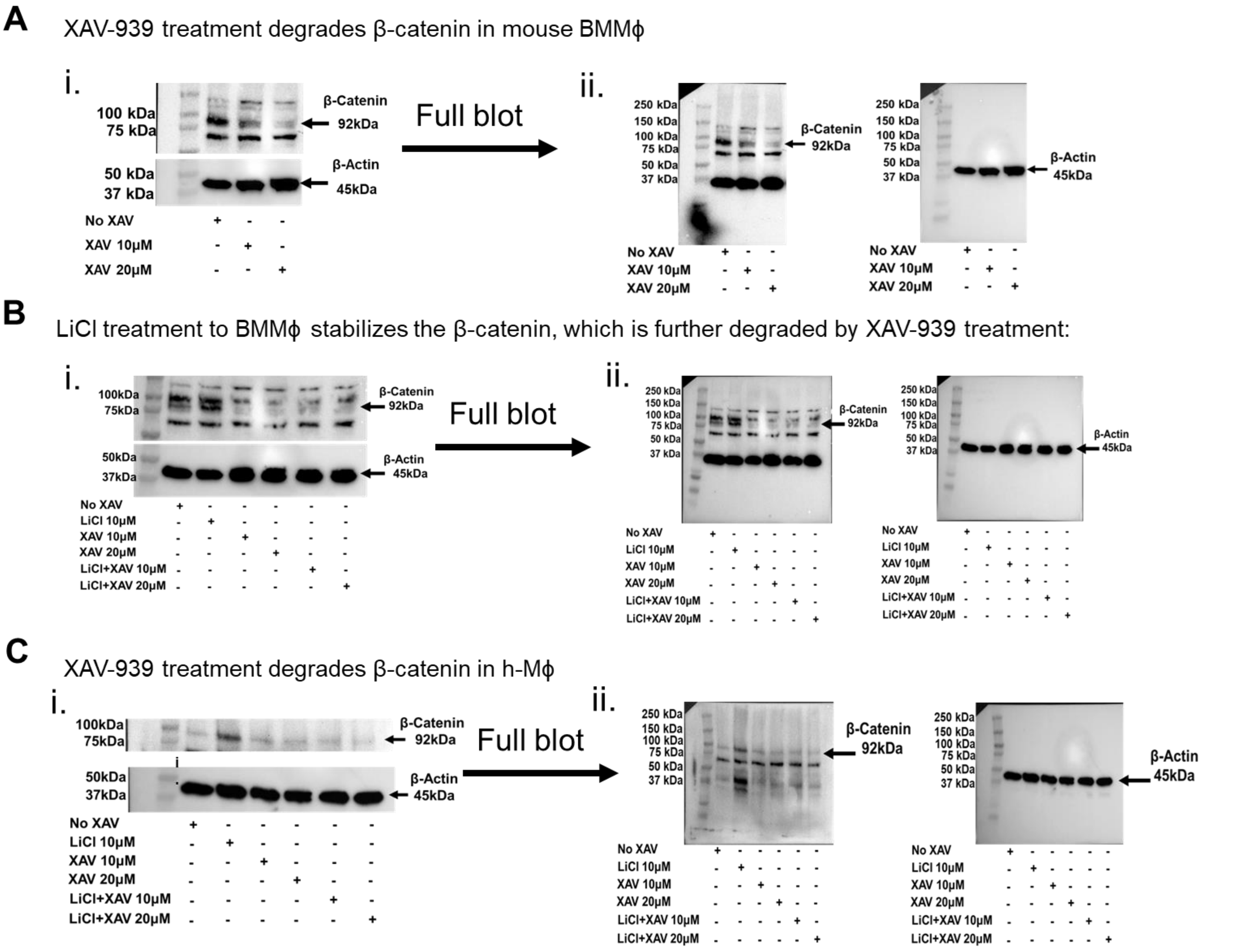


**Supplementary Figure 2: Pharmacological degradation of β-Catenin using XAV-939:** (A i) Immunoblot to detect β-catenin degradation using different concentration of XAV-939 in mouse BMMϕ, (A ii) uncropped Immunoblot of A i. (B i) Immunoblot to detect LiCl treatment stabilizes the β-catenin in BMMϕ, which was further degraded by XAV-939 treatment in BMMϕ pre-treated with LiCl. (B ii) uncropped Immunoblot of B i. (C i) Immunoblot to detect β-catenin degradation using different concentration of XAV-939 in human Mϕ. (C ii) uncropped Immunoblot of C i.

**Supplementary Figure 3:**

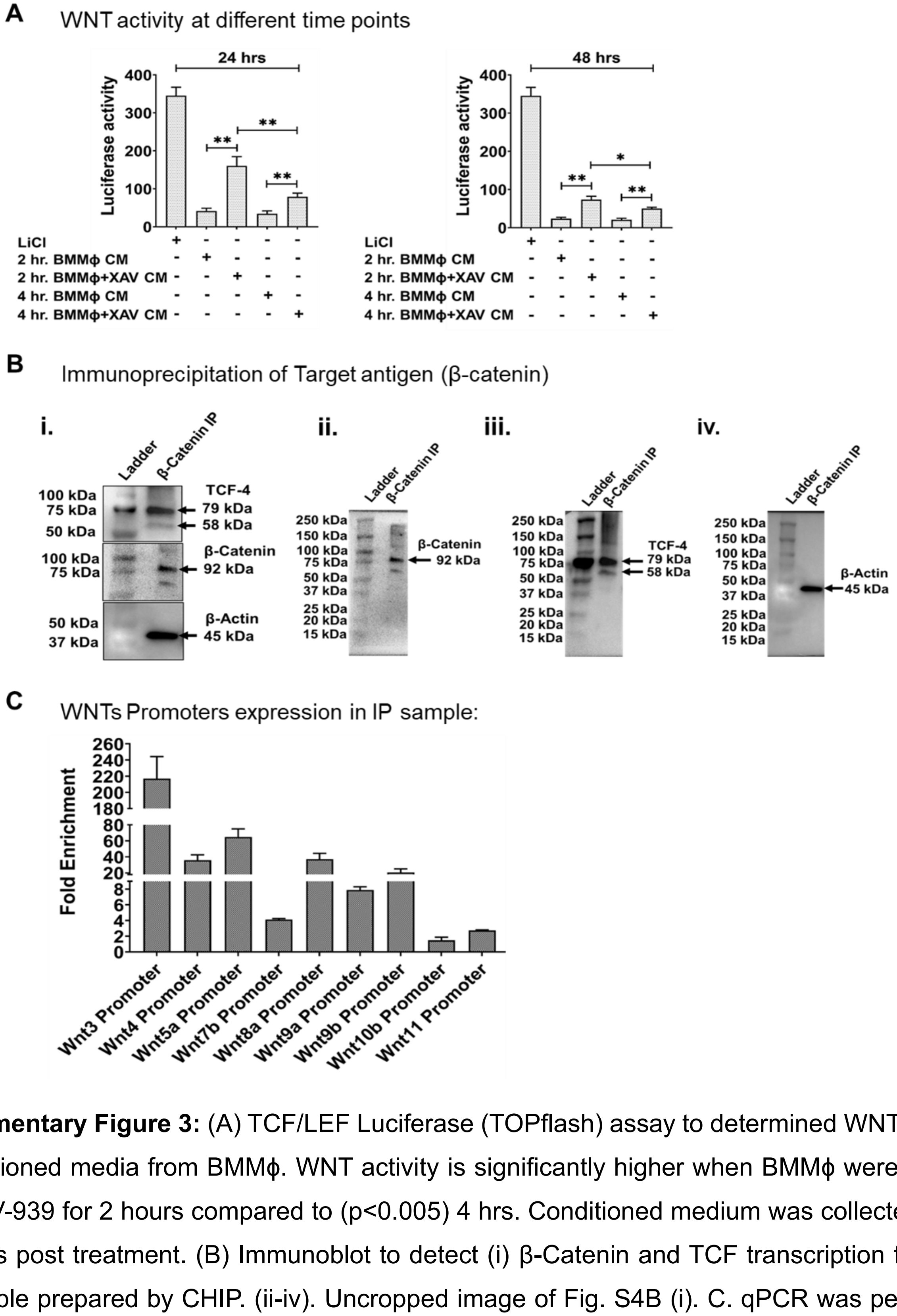


**Supplementary Figure 3:** (A) TCF/LEF Luciferase (TOPflash) assay to determined WNT activity in conditioned media from BMMϕ. WNT activity is significantly higher when BMMϕ were treated with XAV-939 for 2 hours compared to ($p<0.005$) 4 hrs. Conditioned medium was collected at 24 or 48 hrs post treatment. (B) Immunoblot to detect (i) β-Catenin and TCF transcription factor in the sample prepared by CHIP. (ii-iv). Uncropped image of Fig. S4B (i). C. qPCR was performed on the Chip DNA sample to confirm the presence of various WNT promoters using WNT promoter specific primers. Data presented as the mean ± SD. (Significant level, *: $p<0.05$, **: $p<0.005$).

### Supplementary Figure 4:

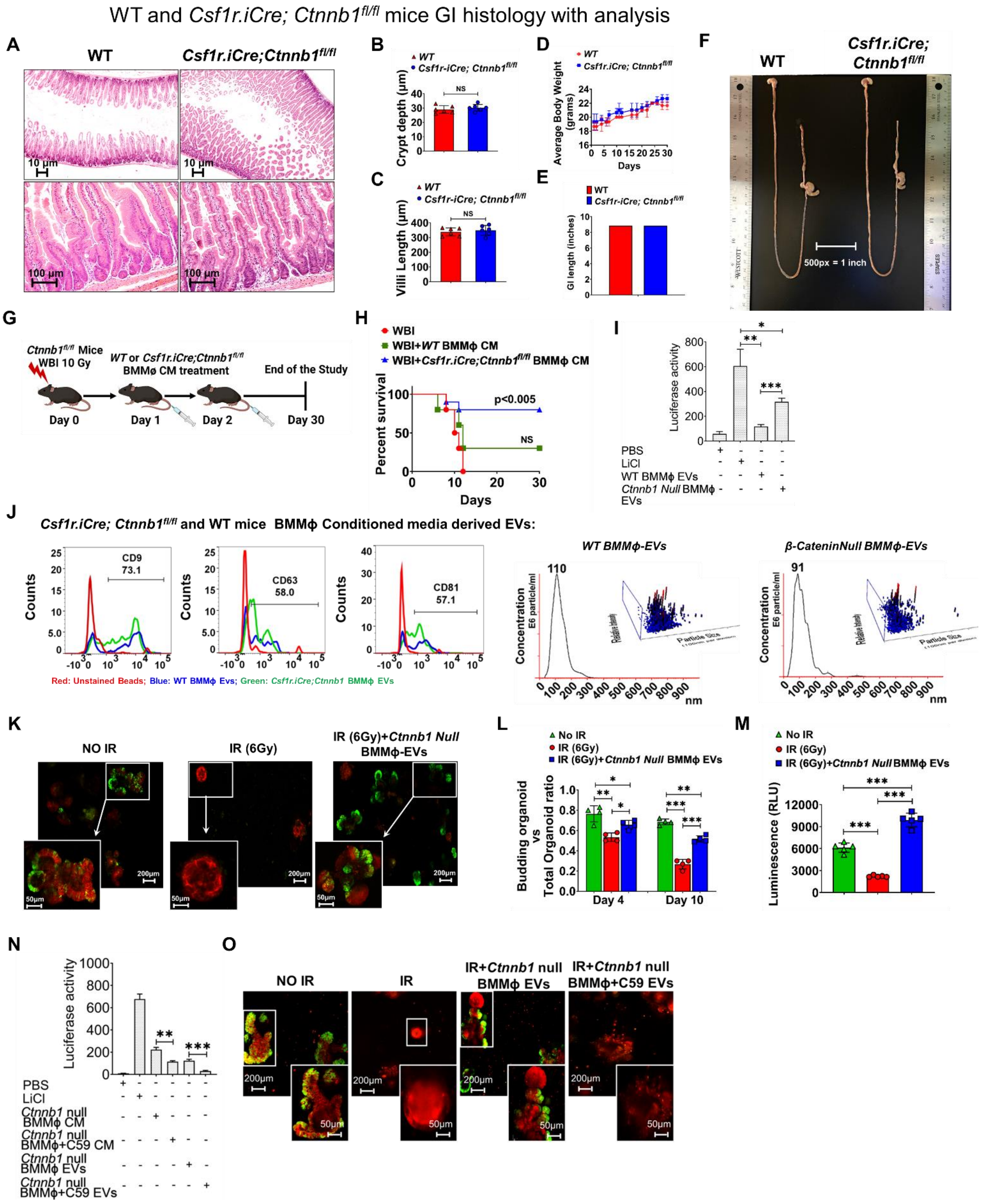

**Supplementary Figure 4: Genetic ablation of β-Catenin in macrophages did not affect mice intestinal development**: (A-C) Hematoxylin and Eosin staining of jejunum section from WT and *Csf1r.iCre; Ctnnb1*$^{fl/fl}$ mice showed no significant differences in crypt depth (A, B), and Villi length (A, C). No differences in (D) average body weight and (E, F) total length of the small intestine was noted between WT and *Csf1r.iCre; Ctnnb1*$^{fl/fl}$ mice. (G) Schematic representation of survival experimental plan (radiation doses and timeline for survival analysis) (H) Kaplan–Meier survival (Mantel-Cox test) analysis of *WT* mice exposed to whole-body irradiation (10 Gy) shows 100% lethality within 10-12 days post irradiation. Significant (80%) (p<0.005) improvement in survival was noted in WT mice receiving *Csf1r.iCre; Ctnnb1*$^{fl/fl}$ BMMϕ conditioned media. Only 30% survival was noted in WT mice receiving WT BMMϕ conditioned media. (I) EVs from *Csf1r.iCre; Ctnnb1*$^{fl/fl}$ BMMϕ conditioned media showed higher luciferase activity in TOPflash assay compared to WT BMMϕ conditioned media EVs. (J) shows *Csf1r.iCre; Ctnnb1*$^{fl/fl}$ and WT BMMϕ EVs characterization by flowcytometric analysis show the presence of exosomal marker CD9 (73.1%) CD63 (58%) CD81 (57.1%) respectively. and EVs particle counts estimated by Nanoparticle tracking analysis (NTA). (K) Confocal microscopic images of organoids developed from Lgr5-EGFP-CRE-ERT2; R26- ACTB-tdTomato-EGFP mice demonstrated that *Csf1r.iCre; Ctnnb1*$^{fl/fl}$ BMMϕ conditioned media derived EV treatment increases the presence of Lgr5-positive cells (green) in budding crypts compared with irradiated controls. tdTomato is constitutively expressed in these mice as membrane bound protein, and therefore allows better visualization of cellular morphology, (L) Histogram demonstrating the significant improvement in irradiated crypt organoid growth following treatment with EVs derived from *Csf1r.iCre; Ctnnb1*$^{fl/fl}$ BMMϕ conditioned media compared to irradiated untreated organoid on day 4 and day 10 post irradiation. (M) Histogram demonstrating significant increase in relative fluorescence units (RFU) immune-fluorescence signal in irradiated intestinal organoids treated with/without *Csf1r.iCre; Ctnnb1*$^{fl/fl}$ BMMϕ conditioned media derived EV. RFU represented the survival of the irradiated organoids. (N) TOPflash assay results demonstrating reduced WNT activity in C59 treated BMMϕ conditioned media (CM) or conditioned media derived EVs. (O) Confocal microscopic images of crypt organoids from Lgr5-EGFP-CRE-ERT2; R26- ACTB-tdTomato-EGFP mice. Organoids treated with C59 treated BMMϕ CM derived EVs of *Csf1r.iCre; Ctnnb1*$^{fl/fl}$ mice did not rescued the LGR5 +ve (GFP +ve cells) ISCs from radiation injury. Data presented as the mean ± SD. (Significant level, *: p<0.05, **: p<0.005, ***: p<0.0005).

**Supplementary Figure 5:**

**A**

**C57BL/6 BMMϕ+XAV v/s C57BL/6 BMMϕ**

| Gene | FC | P Value | FDR |
|---|---|---|---|
| **M1 macrophage markers** | | | |
| Ccl8 | -2.288 | 4.02E-01 | 0.881 |
| Ccl4 | -1.196 | 1.13E-01 | 0.612 |
| Ccl9 | -1.711 | 4.05E-07 | 0.000 |
| Il1b | -2.015 | 1.21E-01 | NA |
| Ccl2 | -1.151 | 2.57E-01 | 0.799 |
| Tlr4 | -1.184 | 5.34E-02 | 0.454 |
| Socs3 | -1.081 | 8.60E-01 | 0.984 |
| **M2 macrophage markers** | | | |
| Il4 | 1.222 | 4.23E-01 | 0.891 |
| Tgfb1 | 1.013 | 7.97E-01 | 0.975 |
| Arg1 | 1.166 | 8.29E-01 | 0.982 |
| Cd200 | 1.401 | 1.42E-01 | 0.664 |
| Cd200r1 | 1.066 | 3.18E-01 | 0.839 |
| Ccl22 | 1.041 | 9.19E-01 | 0.991 |
| **Growth factor** | | | |
| Manf | 1.414 | 8.51E-03 | 0.167 |
| Cxcl1 | 1.741 | 1.33E-02 | 0.218 |
| Vegfb | 1.246 | 2.82E-02 | 0.330 |
| **Anti-Inflammatory** | | | |
| Gene Name | FC | P Value | FDR |
| Cd24a | 2.505 | 6.57E-03 | 0.147 |
| Ier3 | 1.513 | 7.00E-03 | 0.151 |
| Ppard | 1.259 | 1.50E-02 | 0.232 |
| Rora | 1.200 | 2.62E-02 | 0.319 |
| Vps35 | 1.127 | 5.15E-02 | 0.450 |
| **Inflammatory Genes** | | | |
| Gene Name | FC | P Value | FDR |
| Ldlr | -2.651 | 3.21E-21 | 0.000 |
| Fcgr1 | -2.078 | 5.27E-12 | 0.000 |
| Grn | -1.212 | 1.28E-04 | 0.008 |
| Ctsc | -1.501 | 5.78E-04 | 0.028 |
| Zbp1 | -1.769 | 6.74E-04 | 0.031 |
| Alox5ap | -1.208 | 7.33E-04 | 0.033 |
| Stap1 | -1.413 | 1.07E-03 | 0.043 |
| Nfkbiz | -1.247 | 1.20E-03 | 0.048 |
| Tnfrsf11a | -1.329 | 1.91E-03 | 0.066 |
| Tlr9 | -1.936 | 3.99E-03 | 0.104 |
| Ccr2 | -1.929 | 7.27E-03 | 0.156 |
| Ccr5 | -1.323 | 7.71E-03 | 0.160 |
| Ets1 | -1.348 | 9.77E-03 | 0.184 |
| Ifi35 | -1.386 | 1.55E-02 | 0.236 |
| Cd81 | -1.119 | 2.21E-02 | 0.291 |
| Ptger4 | -1.285 | 2.32E-02 | 0.300 |
| Btk | -1.166 | 2.34E-02 | 0.300 |
| Cd47 | -1.137 | 2.96E-02 | 0.338 |
| Tlr3 | -1.224 | 3.85E-02 | 0.387 |
| Tlr4 | -1.184 | 5.34E-02 | 0.454 |

**B**

***Csf1r.iCre;Ctnnb1$^{fl/fl}$* vs WT**

| Gene | FC | P Value | FDR |
|---|---|---|---|
| **M1 macrophage markers** | | | |
| Ccl3 | -1.551 | 4.57E-08 | 0.000 |
| Ccl8 | -3.876 | 1.70E-01 | 0.844 |
| Ccl4 | -1.688 | 3.94E-06 | 0.001 |
| Ccl9 | -1.128 | 2.53E-01 | 0.904 |
| Cd86 | -1.310 | 5.56E-03 | 0.239 |
| Tnf | -1.205 | 1.53E-01 | 0.834 |
| Il1r1 | -2.135 | 2.57E-02 | 0.487 |
| Il1b | -2.544 | 4.17E-02 | 0.598 |
| Ccl2 | -1.620 | 9.71E-05 | 0.014 |
| Il23r | -1.716 | 4.84E-01 | 0.982 |
| Cd80 | -1.115 | 3.20E-01 | 0.937 |
| **M2 macrophage markers** | | | |
| Il4 | 1.136 | 6.11E-01 | 0.995 |
| Tgfb1 | 1.088 | 8.28E-02 | 0.727 |
| Cd163 | 1.192 | 5.56E-01 | 0.992 |
| Arg1 | 1.963 | 3.43E-01 | 0.947 |
| Cd200 | 1.071 | 7.67E-01 | 0.998 |
| Cd200r1 | 1.088 | 1.87E-01 | 0.863 |
| **Growth factor** | | | |
| Gpi1 | 1.162 | 1.04E-02 | 0.331 |
| Gmfg | 1.212 | 1.22E-02 | 0.359 |
| Igf1 | 1.094 | 8.21E-02 | 0.727 |
| Tgfb1 | 1.088 | 8.28E-02 | 0.727 |
| Fgf2 | 1.508 | 8.37E-02 | 0.729 |
| **Anti-Inflammatory** | | | |
| Gene | FC | P Value | FDR |
| Gps2 | 1.235 | 6.81E-03 | 0.267 |
| Lpcat3 | 1.232 | 1.02E-02 | 0.331 |
| Ldlr | 1.279 | 1.62E-02 | 0.411 |
| Selenos | 1.235 | 3.83E-02 | 0.581 |
| **Inflammatory Genes** | | | |
| Gene | FC | P Value | FDR |
| Ccl3 | -1.551 | 4.57E-08 | 0.000 |
| Gprc5b | -1.455 | 6.98E-05 | 0.011 |
| Ptgs2 | -2.058 | 5.09E-03 | 0.229 |
| Napepld | -1.290 | 1.93E-02 | 0.440 |
| Nfkbiz | -1.172 | 1.97E-02 | 0.443 |
| Ets1 | -1.301 | 2.13E-02 | 0.459 |
| Rps19 | -1.138 | 2.86E-02 | 0.506 |
| Il1b | -2.544 | 4.17E-02 | 0.598 |

**Supplementary Figure 5: RNA Seq analysis: (A)** RNA seq analysis of BMMϕ revealed significant upregulation of M2 macrophage, growth factors and anti-inflammatory genes while significant downregulation of the inflammatory genes in pre-conditioned BMMϕ (C57BL/6

BMMϕ+XAV) compared to the untreated C57BL/6 BMMϕ. **(B)** Similarly, RNA seq analysis of BMMϕ from *Csf1r.iCre; Ctnnb1*$^{fl/fl}$ mice showed significant increase in expression of M2 macrophage, growth factors and anti-inflammatory gene while significantly downregulation of the inflammatory gene expression compared to the WT BMMϕ.

### Supplementary Figure 6

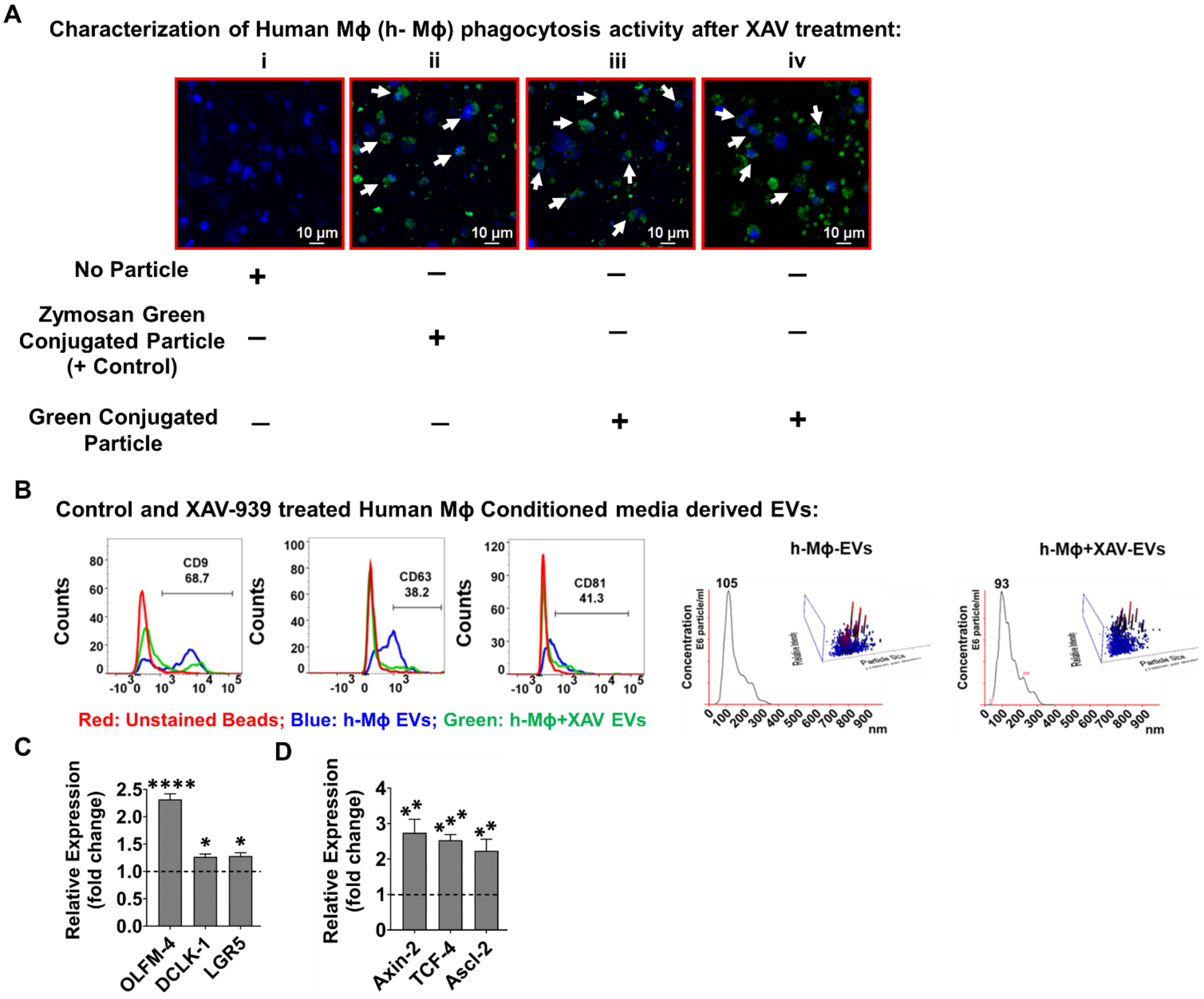


**Supplementary Figure 6: Pharmacological inhibition of β-Catenin in human macrophages (h-Mϕ) does not alter its activity**: (A) Phagocytic activity of Human macrophages was analyzed with or without treatment with XAV-939 using the pHrodo™ Green E. coli BioParticles™ or pHrodo Green Zymosan A BioParticles™. The confocal microscopic images of the human macrophages treated with XAV-939 show no changes in the phagocytic activity suggesting that pharmacological degradation of β-Catenin in human macrophages does not alter its phagocytic characteristic. (B) shows human macrophage (h-Mϕ) and pre-conditioned h-Mϕ EVs characterization by flowcytometric analysis show the presence of exosomal marker CD9 (68.7%) CD63 (38.2%) CD81 (41.3%) respectively and EVs particle counts estimated by Nanoparticle tracking analysis (NTA). (C) shows expression of intestinal stem cells markers OLFM-4 ($p<0.00005$), DCLK-1 ($p<0.05$), LGR5 ($p<0.05$) and (D) β-catenin target genes Axin-2 ($p<0.005$), TCF-4 ($p<0.0005$), AsCl-2 ($p<0.005$), expression in irradiated human intestinal organoid followed by treatment with pre-conditioned h-Mϕ EVs in comparison to the no EVs treatment control. Data presented as the mean ± SD. (Significant level, *: $p<0.05$, **: $p<0.005$, ***: $p<0.0005$, ****: $p<0.00005$).

**Supplementary Figure 7:**

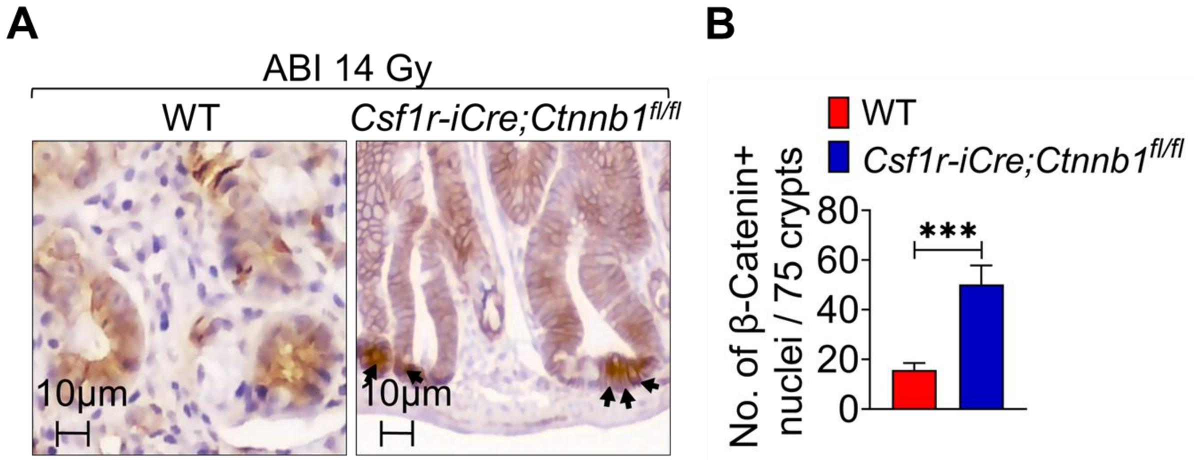


**Supplementary Figure 7:** ***Csf1r.iCre; Ctnnb1**$^{fl/fl}$* **mice with WNT enriched mucosal macrophages are resistant to RIGS:** (A) Representative microscopic images (x40 magnification) of jejunal sections immune-stained with anti β-Catenin antibody to determine β-Catenin nuclear localization. Nucleus stained with hematoxylin. Csf1r.iCre; Ctnnb1fl/fl mice showed more nuclear β-Catenin staining (dark brown; indicated with arrows), compared to WT mice following 14 Gy of abdominal irradiation (ABI). (B) Nuclear β-Catenin count: each data point is the average of the number of β-Catenin-positive nucleus from 15 crypts per field, 5 fields per mice. Number of β-Catenin-positive nucleus in irradiated Csf1r.iCre; Ctnnb1fl/fl mice is significantly higher compared to WT mice Data presented as the mean ± SD. (Significant level; ***: $p<0.0005$).

**Supplementary Figure 8:**

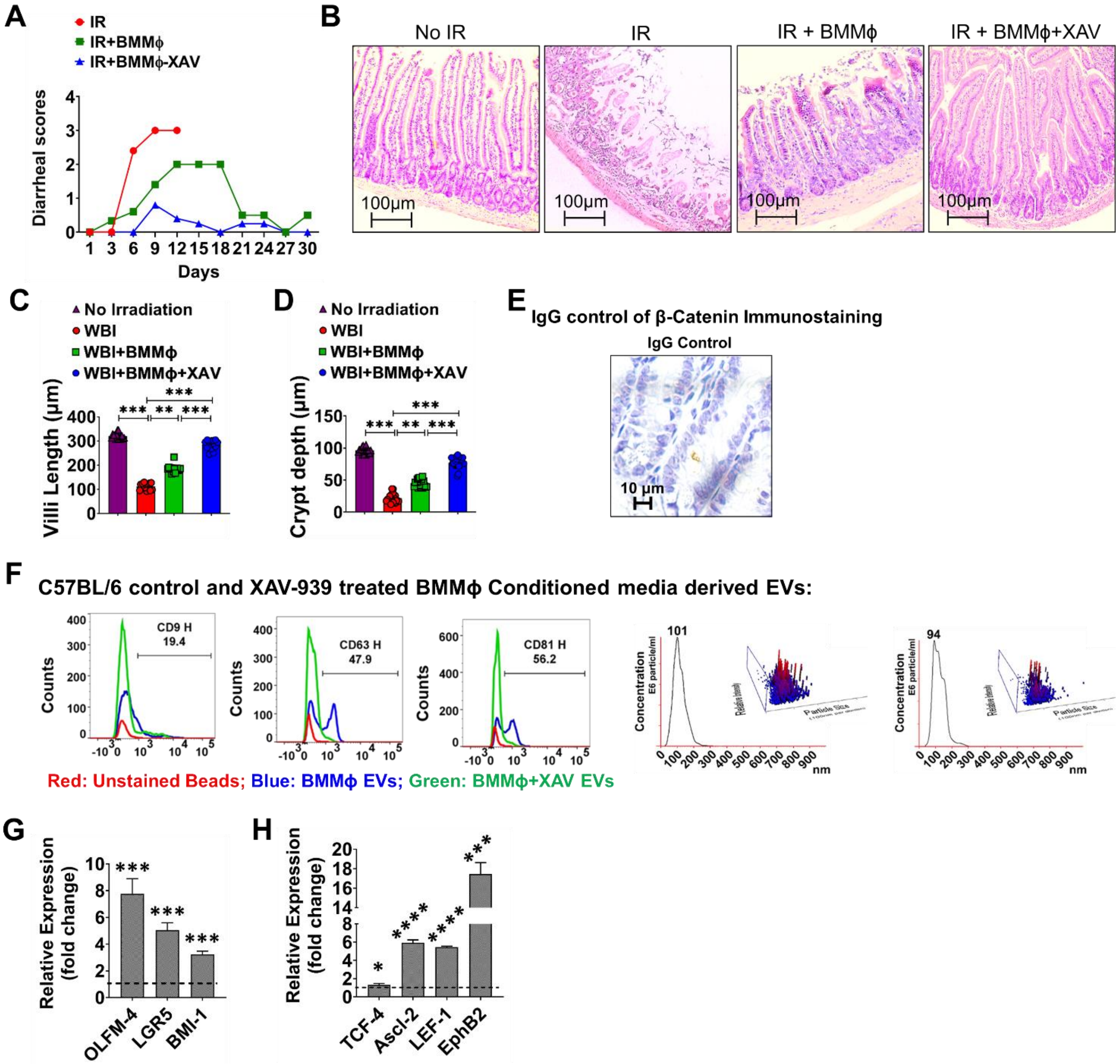


**Supplementary Figure 8:** (A) BMMϕ cell-therapy protect mice from RIGS: C57BL/6 mice receiving BMMϕ, or pre-conditioned BMMϕ post irradiation shows lower diarrheal score than the irradiated control mice (Diarrheal score: 4: watery black stool, 3: Stool with unshaped leave wet mark on the paper towel 2: Normal shape stool but leaves wet stain on paper towel, 1: Normal stool no stain or mark on the paper towel). (B) Hematoxylin and Eosin staining of jejunum section from mice on day 7 after exposed to 10 Gy whole-body irradiation followed by treatment with either BMMϕ or pre-conditioned BMMϕ. (C-D). Histogram showing villus length and crypt depth in jejunal section of C57BL6 mice. Irradiated C57BL6 mice demonstrated a significant decrease in villi length and crypt depth. Irradiated C57BL6 mice receiving pre-conditioned BMMϕ showing less reduction of villi length and crypt depth compared to irradiated mice receiving native BMMϕ or no cell therapy. (E) Immunostaining with IgG control for β-Catenin antibody. (F) C57BL/6 BMMϕ and pre-conditioned BMMϕ EVs characterization by flowcytometric analysis shows the presence of exosomal marker CD9 (19.4%) CD63 (47.9%) CD81 (56.2%) respectively and EVs particle counts estimated by Nanoparticle tracking analysis (NTA). (G) shows significant increased expression of intestinal stem cells markers in intestinal stem cells marker OLFM-4 ($p<0.0005$), LGR5 ($p<0.0005$), BMI-1 ($p<0.0005$) and (H) β-Catenin target genes (TCF-4 ($p<0.05$), AsCl-2 ($p<0.00005$), LEF-1 ($p<0.00005$), EphB2 ($p<0.0005$) expression was noted in irradiated mouse intestinal organoid in response to pre-conditioned BMMϕ EV treatment compared to untreated irradiated control. Data presented as the mean ± SD. (Significant level, *: $p<0.05$, **: $p<0.005$, ***: $p<0.0005$, ****: $p<0.00005$).

**Supplementary Figure 9:**

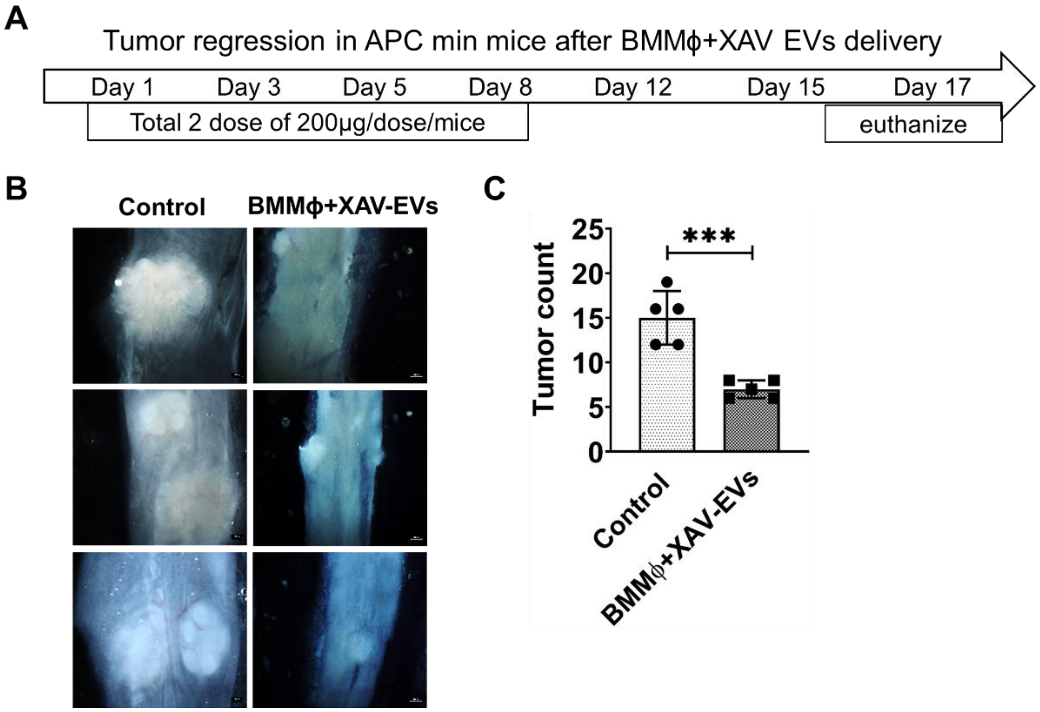


**Supplementary Figure 9: Pre-conditioned BMMϕ EVs modulate Intestinal tumor growth: (A)** Experimental plan for the treatment of APC min+ mice with/without pre-conditioned BMMϕ EVs for tumor regression analysis. **(B)** Representative images of the tumors in control and Pre-conditioned BMMϕ EVs treated mice. **(C)** Histogram demonstrating tumor number. Number of tumor /mice was significantly lower in Pre-conditioned BMMϕ EVs treated mice compared to the control untreated mice. (n=3 mice per group) Data presented as the mean ± SD. (Significant level, ***: $p<0.0005$).

**Supplementary Table 1: Identified Wnt promoters based on their confidence score.**

| Chromosome | Peak start | Peak end | Confidence Score | Identified Promoter |
|---|---|---|---|---|
| chr11 | 59306380 | 59306428 | 3.789 | Wnt9b_up2k |
| chr11 | 59305712 | 59305760 | 2.315 | Wnt9b_up2k |
| chr14 | 28510318 | 28510366 | 2.315 | Wnt5a_up2k |
| chr11 | 59291109 | 59291157 | 2.284 | Wnt3a_up2k |
| chr11 | 103773730 | 103773778 | 2.180 | Wnt3_up2k |
| chr6 | 18032018 | 18032066 | 1.876 | Wnt2_up2k |
| chr1 | 74769488 | 74771335 | 1.872 | Wnt6_up2k |
| chr15 | 85579495 | 85580058 | 1.809 | Wnt7b_up2k |
| chr11 | 103772404 | 103772474 | 1.498 | Wnt3_up2k |
| chr4 | 137276159 | 137276902 | 1.481 | Wnt4_up2k |
| chr7 | 98837090 | 98837319 | 1.448 | Wnt11_up2k |
| chr19 | 44491977 | 44492050 | 1.394 | Wnt8b_up2k |
| chr4 | 137275598 | 137275840 | 1.386 | Wnt4_up2k |
| chr15 | 85581166 | 85581663 | 1.340 | Wnt7b_up2k |
| chr1 | 74791558 | 74792232 | 1.319 | Wnt10a_up2k |
| chr6 | 91412617 | 91413105 | 1.306 | Wnt7a_up2k |
| chr6 | 22287003 | 22287483 | 1.286 | Wnt16_up2k |
| chr6 | 18030957 | 18031005 | 1.277 | Wnt2_up2k |
| chr6 | 119449325 | 119449797 | 1.246 | Wnt5b_up2k |
| chr7 | 98838342 | 98838467 | 1.245 | Wnt11_up2k |
| chr4 | 137277118 | 137277328 | 1.216 | Wnt4_up2k |
| chr6 | 18031484 | 18031742 | 1.215 | Wnt2_up2k |
| chr6 | 22286150 | 22286483 | 1.193 | Wnt16_up2k |
| chr15 | 98778294 | 98778955 | 1.189 | Wnt10b_up2k |

**Supplementary Table 2:**

**Table S2: Parameter included for analyzing DSS induced Colitis in mice:**

**A. Disease activity scores (DAI):**

| Parameters | Score |
|---|---|
| Stool consistency | 0 (normal), 2 (loose stool), and 4 (diarrhea); |
| Bleeding | 0 (no blood), 1 (Hemoccult positive), 2 (Hemoccult positive and visual pellet bleeding), and 4 (gross bleeding, blood around anus). |
| % Weight loss | 0 (no loss), 1 (1-5%), 2 (5-10%), 3 (10-20%), and 4 (>20%); |

**B. Histology scores:**

| Score | Goblet cell loss | Submucosal inflammation | Crypt density | Inflammatory infiltrate |
|---|---|---|---|---|
| **0** | none | none | normal | none |
| **1** | <10% | individual cells | decreased by <10% | increased presence of inflammatory cells |
| **2** | 10–50% | infiltrate(s) | decreased by 10–50% | infiltrates also in submucosa |
| **3** | >50% | large infiltrate(s) | decreased by >50% | transmural |

**Supplementary Table 3:**

**Primer Sequences:**

**Mouse WNTs Primer Sequence:**

| | Forward Sequence (5'-3') | Reverse Sequence (5'-3') |
|---|---|---|
| Wnt-1 | TCTTTGGCCGAGAGTTCGTG | AGAGAACACGGTCGTTCGC |
| Wnt-2 | ATCTCTTCAGCTGGCGTTGT | AGCCAGCATGTCCTCAGAGT |
| Wnt-2b | CACCCGGACTGATCTTGTCT | TGTTTCTGCACTCCTTGCAC |
| Wnt-3 | TGGAACTGTACCACCATAGATGAG | ACACCAGCCGAGGCGATG |
| Wnt-3a | ACCGTCACAACAATGAGGCT | TCGGCACCTTGAAGTACGTG |
| Wnt-4 | AACGGAACCTTGAGGTGATG | GGACGTCCACAAAGGACTGT |
| Wnt-5a | CACGCTATACCAACTCCTCTGC | AATATTCCAATGGGCTTCTTCATGGC |
| Wnt-5b | GCCGCGGATGAGGAGTG | GCCTCAACCCATCCCAATGC |
| Wnt-6 | CGGAGACGATGTGGACTT | GGAACCCGAAAGCCCATG |
| Wnt-7a | ATCAAGCAGAATGCCCGGAC | TAGCTCTCGGAACTGTGGCA |
| Wnt-7b | ACTCCGAGTAGGGAGTCGAGA | GCGACGAGAAAAGTCGATGC |
| Wnt-8a | TGGGAACGGTGGAATTGTCC | GCAGAGCGGATGGCATGAAT |
| Wnt-8b | GTGGACTTCGAAGCGCTAAC | TTACACGTGCGTTTCATGGT |
| Wnt-9a | TGCTTTCCTCTACGCCATCT | TATCACCTTCACACCCACGA |
| Wnt-9b | GTGTGGTGACAATCTGAAG | TCCAACAGGTACGAACAG |
| Wnt-10a | GCTTCGGAGAACGCTTCTCT | ATTTGCACTTACGCCGCATG |
| Wnt-10b | GGAAGGGTAGTGGTGAGCAA | CACTTCCGCTTCAGGTTTTC |
| Wnt-11 | GTTCTCCGTGATTGCAGGCG | TTGCGTCTGATTCAGTGCCA |
| Wnt-16 | CTGTGACACCACCTTGCAGA | CAGGTTTTCACAGCACAGGA |
| GAPDH | ACCACAGTCCATGCCATCAC | TCCACCACCCTGTTGCTGTA |

**Human WNTs Primer Sequence:**

| | Forward Sequence (5'-3') | Reverse Sequence (5'-3') |
|---|---|---|
| Wnt-1 | GCGTCTGATACGCCAAAATC | GGATTCGATGGAACCTTCTG |
| Wnt-2 | TAGTCGGGAATCTGCCTTTG | TTCCTTTCCTTTGCATCCAC |
| Wnt-2b | CTCATCAGCAGGGGTAGTCC | AAAACGGACACCGTAGTGGA |
| Wnt-3 | ACGAGAACTCCCCCAACTTT | GATGCAGTGGCATTTTTCCT |
| Wnt-3a | TGTTGGGCCACAGTATTCCT | ATGAGCGTGTCACTGCAAAG |
| Wnt-4 | CCTTCGTGTACGCCATCTCT | GCCTCATTGTTGTGGAGGTT |
| Wnt-5a | CCCATGCAGTACATCGGAG | CACTCTCGTAGGAGCCCTTG |
| Wnt-5b | GTGCAGAGACCCGAGATGTT | CAGGCTACGTCTGCCATCTT |
| Wnt-6 | GGTTATGGACCCTACCAGC | AATGTCCTGTTGCAGGATGC |
| Wnt-7a | AGTACAACGAGGCCGTTCAC | GCACGTGTTGCACTTGACAT |
| Wnt-7b | AAGCTCGGAGCACTGTCATC | CCCTCGGCTTGGTTGTAGTA |
| Wnt-8a | TGGGGAACCTGTTTATGCTC | CCCTCGGCTTGGTTGTAGTA |
| Wnt-8b | CTGGTCCAAAGGCTTACCTG | TGAGTGCTGCGTGGACTTC |
| Wnt-9a | GACGGTCAAGCAAGGATCTG | TGCTCTCGCAGTTCTTCTCA |
| Wnt-9b | CTGCTTGAGTGCCAGTTTCA | CGAGTCATAGCGCAGTTTCA |
| Wnt-10a | AATGCCAACACCAATTCAGG | CAACTCGGTTGTTGTGAAGC |
| Wnt-10b | GCAAGAGTTTCCCCCACTCT | GATTGCGGTTGTGGGTATC |
| Wnt-11 | TTG CTT GAC CTG GAG AGA GG | GAC GAG TTC CGA GTC CTT CA |
| Beta-Actin | TGGATCAGCAAGCAGGAGTATG | GCATTTGCGGTGGACGAT |

**Mouse WNT Promoters Primer Sequence:**

| Forward Sequence (5'-3') | Reverse Sequence (5'-3') |
|---|---|

| | | |
|---|---|---|
| Wnt3 Promoter | TTCCGTCTCAGTGTCTCTATCC | CAGATTCACCCTACGACCTACT |
| Wnt4 Promoter | CCTGACGTGGGAGAAGTAATAAA | GAATCCGAAACCTCGCTTCT |
| Wnt5a Promoter | GTGTGTCTGTGTGTGTCTGT | CCTCTTTCCCTTGGTGTTCTC |
| Wnt7b Promoter | CGGCTGTATCGGTATGTCTTT | TCTCTCTCTCTCTCTCTCTCTCT |
| Wnt8a Promoter | GGTGGTGTCGGTAACAGAAATA | AATCAAAGAGGGACGGTTTAGG |
| Wnt9a Promoter | AGG TCT CCC ATC CAG AAT CA | CTGTAGCTGGAGGAATGGAAAG |
| Wnt9b Promoter | AGATGTCAGAGAGGAGGTAGT G | CCAAGGTCTGGGTTGAATCTTA |
| Wnt10b Promoter | CCAGCACTGTCTCAGTAACTTC | GAACCTCTGGGCGCTATAATC |
| Wnt11 Promoter | TCTCTCTCTCTCTCTCTCTCT CT | CCACCTCCGAAGCTAGTAATG |

#### Mouse Stem cells markers Primer Sequence:

| | **Forward Sequence (5'-3')** | **Reverse Sequence (5'-3')** |
|---|---|---|
| OLFM4 | GCCACTTTCCAATTTCAC | GAGCCTCTTCTCATACAC |
| LGR5 | AGAGCCTGATACCATCTG CAA AC | TGAAGGTCGTCCACACTGTTGC |
| BMI-1 | ACTACACGCTAATGGACATTGCC | CTCTCCAGCATTCGTCAGTCC A |

#### Human Stem cells markers Primer Sequence:

| | **Forward Sequence (5'-3')** | **Reverse Sequence (5'-3')** |
|---|---|---|
| OLFM4 | GACCAAGCTGAAAGAGTGTGAGG | CCTCTCCAGTTGAGCTGAACCA |
| LGR5 | CCTGCTTGACTTTGAGGAAGACC | CCAGCCATCAAGCAGGTGTTCA |
| DCLK-1 | ACCGATGCCATCAAGCTGGACT | TCCTGGTAACGGAACTTTCC G |

#### Mouse WNT target genes Primer Sequence:

| | **Forward Sequence (5'-3')** | **Reverse Sequence (5'-3')** |
|---|---|---|
| TCF-4 | GGCGTTGGACAGATCACC | GGTGAAGTGTTCATTGCTGTACTG |
| Ascl-2 | CTACTCGYCGGAGGAAAG | ACTAGACAGCAT GGTAAG |
| LEF-1 | AGACACCCTCCAGCTCCTGA | CCTGAATCCACCCGTGATG |
| EphB2 | CGCCATCTATGTCTTCCAGGTG | GATGAGTGGCAACTTCTCCTG G |

#### Human WNT target genes Primer Sequence:

| | **Forward Sequence (5'-3')** | **Reverse Sequence (5'-3')** |
|---|---|---|
| Axin-2 | CAAACTTTCGCCAACCGTGGTTG | GGTGCAAAGACATAGCCAGAACC |
| TCF-4 | GCCTCTTCACAGTAGCATG | GCTGGTTTGGAGGAAGGATAGC |
| Ascl-2 | CGCCTACTCGTCGGACGACAG | GCCGCTCGCTCGGCTTCCG |
